# Evolution of neuronal cell classes and types in the vertebrate retina

**DOI:** 10.1101/2023.04.07.536039

**Authors:** Joshua Hahn, Aboozar Monavarfeshani, Mu Qiao, Allison Kao, Yvonne Kölsch, Ayush Kumar, Vincent P Kunze, Ashley M. Rasys, Rose Richardson, Herwig Baier, Robert J. Lucas, Wei Li, Markus Meister, Joshua T. Trachtenberg, Wenjun Yan, Yi-Rong Peng, Joshua R. Sanes, Karthik Shekhar

**Affiliations:** Department of Chemical and Biomolecular Engineering, University of California, Berkeley, Berkeley, CA 94720, USA; Department of Cellular and Molecular Biology, Center for Brain Science, Harvard University, MA 02138, USA; Division of Biology and Biological Engineering, California Institute of Technology, Pasadena, CA 91125, USA; Max Planck Institute for Biological Intelligence, Am Klopferspitz 18, 82152 Martinsried, Germany; Retinal Neurophysiology Section, National Eye Institute, National Institutes of Health, Bethesda, MD 20892, USA; Department of Cellular Biology, University of Georgia, Athens, GA 30602; Division of Neuroscience and Centre for Biological Timing, Faculty of Biology Medicine & Health, University of Manchester, Upper Brook Street, Manchester M13 9PT, UK; Department of Neurobiology, David Geffen School of Medicine at UCLA, Los Angeles, CA 90095, USA; Department of Ophthalmology, Stein Eye Institute, UCLA David Geffen School of Medicine, Los Angeles, CA 90095 United States; Helen Wills Neuroscience Institute, Vision Science Graduate Group, Center for Computational Biology, Biophysics Graduate Group, California Institute of Quantitative Biosciences (QB3), University of California, Berkeley, Berkeley CA 94720, USA; Faculty Scientist, Biological Systems and Engineering Division, Lawrence Berkeley National Laboratory, Berkeley, CA 94720, USA

**Author notes:** LinkedIn, Mountain View, CA, 94043. These authors contributed equally to this work (Joshua Hahn, Aboozar Monavarfeshani).

## Abstract

The basic plan of the retina is conserved across vertebrates, yet species differ profoundly in their visual needs (Baden et al., 2020). One might expect that retinal cell types evolved to accommodate these varied needs, but this has not been systematically studied. Here, we generated and integrated single-cell transcriptomic atlases of the retina from 17 species: humans, two non-human primates, four rodents, three ungulates, opossum, ferret, tree shrew, a teleost fish, a bird, a reptile and a lamprey. Molecular conservation of the six retinal cell classes (photoreceptors, horizontal cells, bipolar cells, amacrine cells, retinal ganglion cells [RGCs] and Müller glia) is striking, with transcriptomic differences across species correlated with evolutionary distance. Major subclasses are also conserved, whereas variation among types within classes or subclasses is more pronounced. However, an integrative analysis revealed that numerous types are shared across species based on conserved gene expression programs that likely trace back to the common ancestor of jawed vertebrates. The degree of variation among types increases from the outer retina (photoreceptors) to the inner retina (RGCs), suggesting that evolution acts preferentially to shape the retinal output. Finally, we identified mammalian orthologs of midget RGCs, which comprise >80% of RGCs in the human retina, subserve high-acuity vision, and were believed to be primate-specific (Berson, 2008); in contrast, the mouse orthologs comprise <2% of mouse RGCs. Projections both primate and mouse orthologous types are overrepresented in the thalamus, which supplies the primary visual cortex. We suggest that midget RGCs are not primate innovations, but descendants of evolutionarily ancient types that decreased in size and increased in number as primates evolved, thereby facilitating high visual acuity and increased cortical processing of visual information.

## Main

The ability to assess gene conservation among species has been of great value in multiple ways. It has illuminated the evolutionary history of specific genes, highlighted crucial developmental and functional pathways, informed strategies for rational *in vivo* manipulations, and helped guide choices of animal models that mimic human diseases (Alfoldi and Lindblad-Toh, 2013; Koonin, 2005). Comparative genomics was enabled by advances in DNA sequencing, as well as statistical methodologies for sequence alignment and phylogenetic inference (Durbin et al., 1998). Recent advances in high-throughput single-cell and single-nucleus transcriptomic profiling (scRNA-seq, snRNA-seq) enable a related enterprise focused on determining the extent to which cell types, the functional units of complex tissues (Zeng, 2022; Zeng and Sanes, 2017), are conserved among species. Analyzing patterns of cell type conservation across the species phylogeny can serve as a conceptual foundation for reconstructing the evolution of cell types and identifying conserved developmental programs (Marioni and Arendt, 2017; Tanay and Sebe-Pedros, 2021).

The neural retina, the portion of the brain that resides in the back of the eye, is well-suited for this type of analysis. It is arguably as complex as any other part of the brain, but its compactness and accessibility facilitate detailed investigations of structure and function (Dowling, 2012). Moreover, unlike other brain regions (e.g., the cerebral cortex), the retina’s basic structural blueprint is highly conserved among vertebrates (Baden *et al*., 2020). The retina contains five neuronal classes –photoreceptors (PRs), horizontal cells (HCs), bipolar cells (BCs), amacrine cells (ACs), and retinal ganglion cells (RGCs) – and a resident glial class called Müller glia (MG) (Masland, 2012). The cell somata are arranged in three nuclear layers separated by two plexiform (synaptic) layers (**Fig. 1a**) with information flowing through them in a defined direction: PRs in the outer nuclear layer sense light and transmit visually-evoked signals to interneurons in the inner nuclear layer; the interneurons (HCs, BCs, and ACs) process the information and supply it to RGCs in the innermost layer; and the RGCs send axons through the optic nerve to visual centers in the brain. Importantly, most of the neuronal classes can be subdivided into multiple subclasses, and most subclasses comprise multiple types that differ in morphology, physiology, connectivity and molecular composition (Cajal, 1893; Dowling, 2012; Masland, 2012; Sanes and Masland, 2015; Zeng and Sanes, 2017). The specificity of connections between interneuronal and RGC types endows each RGC type with selective responsiveness to small subsets of visual features such as edges, directional motion, and chromaticity (Kerschensteiner, 2022; Sanes and Masland, 2015). As a result of neural computations in the retina, the optic nerve transmits a set of parallel representations of the visual scene to the rest of the brain for further processing (Martersteck et al., 2017; Robles et al., 2014).

**Fig. 1.**
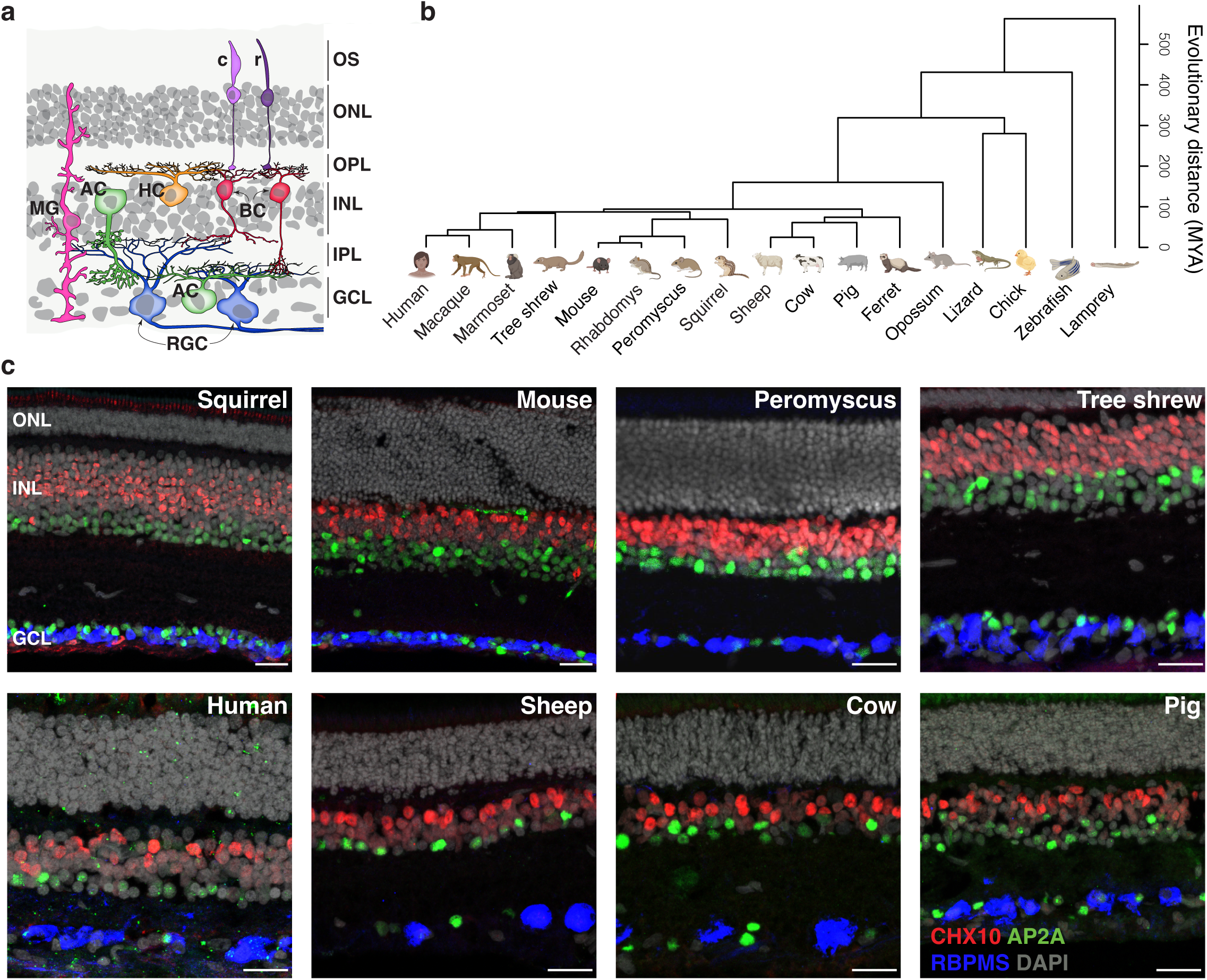
Conserved retinal structure across vertebrates. a. Cartoon of a section through a vertebrate retina showing the arrangement of its six major cell classes – photoreceptors (PRs), horizontal cells (HCs), bipolar cells (BCs), amacrine cells (ACs), retinal ganglion cells (RGCs), and Müller glia (MG). PRs include rods (r) and cones (c). Outer and inner nuclear layers (ONL and INL) and ganglion cell layer (GCL), which contain cell somata, are indicated as are the outer and inner plexiform (synaptic) layers (OPL, IPL). b. Phylogeny of the 17 vertebrate species analyzed in this work. For Linnaean names, see **Supplementary Table 1**. Scale bar on the right indicates estimated divergence time in “millions of years ago” (MYA). c. Sections from retinas of eight species immunostained for RBPMS (a pan-RGC marker), CHX10/VSX2 (a pan-BC marker), AP2A/TFAP2A (a pan-AC marker), and DAPI (a nuclear stain). Colors of the labeled cell classes as in **a.** Scale bars, 25 µm.

Despite these conserved features, vertebrate species differ greatly in their visual needs(Baden *et al*., 2020). Some species are diurnal, others nocturnal; some are terrestrial, others aquatic; some mainly hunt, others forage for colorful fruits. It is likely that variations in retinal cell types across species emerged during the course of evolution to serve these diverse needs. However, a quantitative understanding of evolutionary variation in retinal cell types has been missing. Here, we address this gap by using single-cell transcriptomics to compare retinal cell classes, subclasses and types in 17 vertebrate species (**Fig. 1b,c**).

First, we show that the conserved functional and morphological character of the six cell classes is mirrored by striking cross-species similarities in gene expression. This principle extends to identified subclasses of PRs, BCs, and ACs. Transcription factors implicated in cell and subclass specification are also evolutionarily conserved, pointing to common programs of retinal development. Within each cell class, the transcriptomic variation across species exhibits hallmarks of neutral drift as well as stabilizing selection (Chen et al., 2019). Second, we assessed the extent of evolutionary variation among cell types within BCs and RGCs, two highly diverse retinal classes that have been comprehensively classified in mice (Rheaume et al., 2018; Shekhar et al., 2016; Tran et al., 2019) and primates (Cowan et al., 2020; Peng et al., 2019; Yan et al., 2020b). We identify numerous evolutionarily conserved types but find that variation in RGC types is more extensive than in other classes, suggesting that natural selection acts preferentially to shape the retinal output. Finally, we identify non-primate orthologs of midget RGCs, which account for >80% of RGCs in humans and are primarily responsible for high-acuity vision. To date, no counterparts of these types have been identified in non-primates, precluding mechanistic analysis of blinding diseases involving RGC loss such as glaucoma. This orthology suggests that rather than being evolutionary innovations, midget RGCs descended from types present in the common mammalian ancestor. Surprisingly, murine orthologs of midgets and parasols comprise less than <2% of mouse RGCs (Tran *et al*., 2019). We present evidence that their >20-fold expansion in the primate retina may be related to the increased role of the cerebral cortex in primate visual processing.

## Retinal cell atlases of 17 species

Previously, we used sc/snRNA-seq to study retinal cell types in five species: laboratory mouse (*Mus musculus*, called “mouse” here) (Macosko et al., 2015; Shekhar *et al*., 2016; Tran *et al*., 2019; Yan et al., 2020a), cynomolgus macaque (*Macaca fascicularis*) (Peng *et al*., 2019), human (*Homo sapiens*)(Yan *et al*., 2020b), chick (*Gallus gallus domesticus*) (Yamagata et al., 2021), and zebrafish (*Danio rerio*) (Kolsch et al., 2021) (Yoshimatsu et al., in preparation). For the present study, we generated atlases from twelve additional species: ferret (*Mustela putoriusfuro*), brown anole lizard (*Anolis sagrei*), deer mouse (*Peromyscus maniculatus bairdii*), tree shrew (*Tupaia belangeri chinensis*), pig (*Sus domesticus*), sheep (*Ovis aries*), cow (*Bos taurus*), opossum (*Monodelphis domestica*), marmoset (*Callithrix jacchus*), four-striped grass mouse (*Rhabdomys pumilio*), thirteen-lined ground squirrel (*Ictidomys tridecemlineatus*), and sea lamprey (*Petromyzon marinus*)(**Fig. 1b,c**). We also profiled ∼185,000 nuclei from an additional 18 human donors, thereby allowing us to identify over 30 more types than had been detected in the dataset analyzed previously (Yan *et al*., 2020b), including 10 additional RGC types (**Extended Data Fig. 1**). To obtain sufficient numbers of BCs and RGCs for comprehensive analysis, we enriched these classes in some collections (**Methods**; **Extended Data Fig. 2**). We also collected cells without enrichment to ensure representation of all classes.

**Fig. 2.**
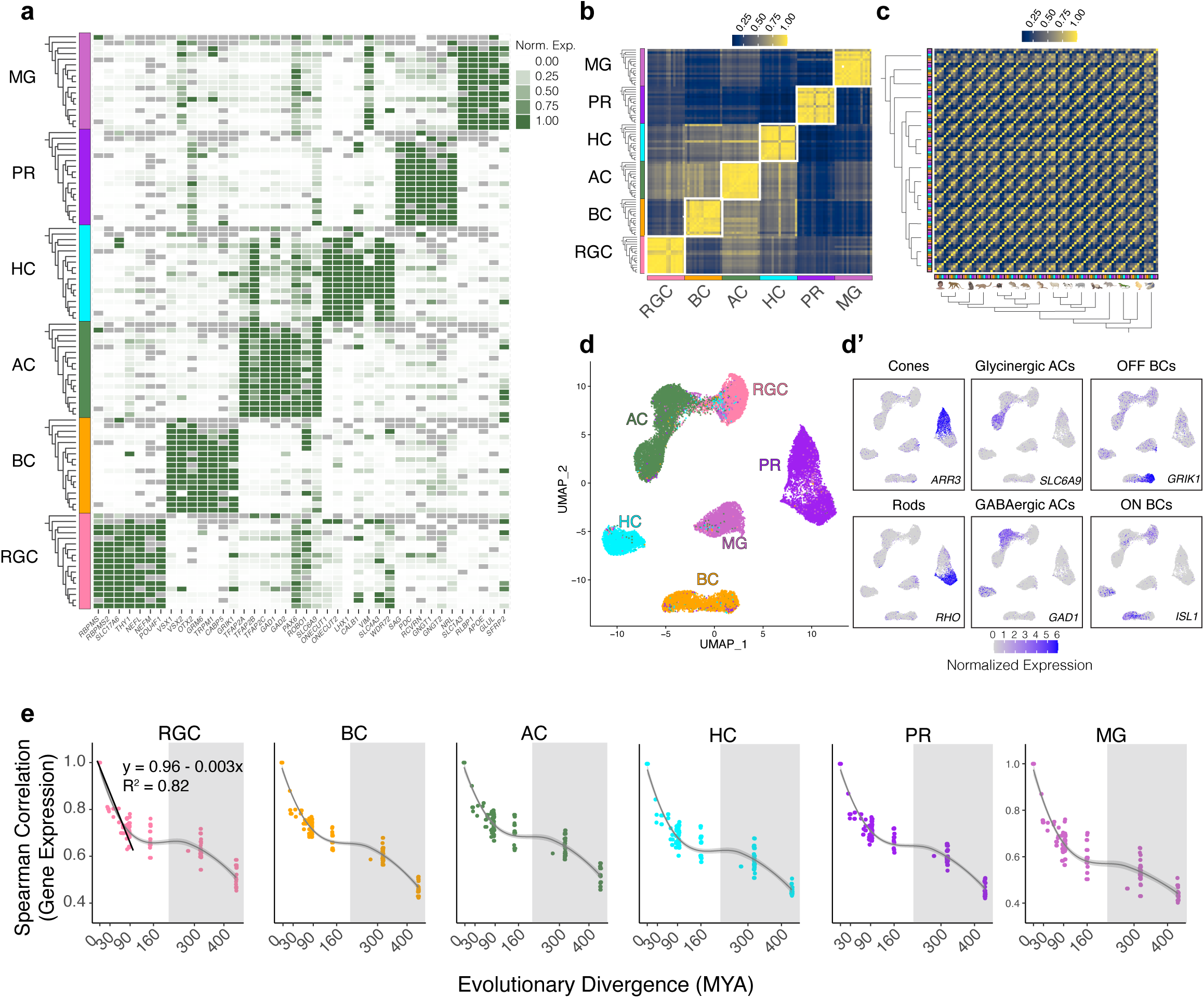
Class- and subclass-specific transcriptomic signatures. a. Heatmap showing average expression of marker genes (columns) within each major cell class in 17 species (rows). Rows are grouped by cell class (annotation bar, left). Within each class, species are ordered as in **Fig. 1b**, with top and bottom nodes in each dendrogram corresponding to lamprey and human, respectively (corresponding to right and left in **Fig. 1b**). Colors indicating cell class are uniform across panels (e.g. RGC is pink). b. Cross-correlation matrix (Spearman) of pseudobulk transcriptomic profiles for the 16 jawed vertebrates. Rows and columns are grouped by class, and then ordered by phylogeny within a class. c. Same as **b**, with rows and columns grouped by species instead of class. Matrices including lamprey (a jawless vertebrate) are shown in **Extended Data Fig. 7c,d**. d. UMAP embedding of integrated cross-species data, with points indicating class identity. d’. Same as **d**, with panels showing cells colored by their expression levels of subclass-specific markers. Within each panel, the gene and labeled cell subclass are indicated. The left, middle and right panel columns correspond to subclasses of PRs (Cones, *upper*; Rods, *lower*), ACs (Glycinergic ACs, *upper*; GABAergic ACs, *lower*) and BCs (OFF BCs, *upper*; ON BCs, *lower*). GAD1, a marker for GABAergic ACs among ACs, is also expressed by some HCs, and *ISL1*, a marker for ON BCs among BCs, is also expressed by some RGCs, HCs, and ACs. Details of gene expression by species are shown in **Extended Data Fig. 8d**. e. Pairwise correlation coefficients of class-specific cell-averaged profiles between species (y-axis) decreases with evolutionary divergence (x-axis) for each cell class. Gray shaded regions demarcate pairwise comparisons involving a non-mammalian vertebrate. In each panel, the trendlines were estimated using locally estimated scatterplot smoothing regression (LOESS). The y vs. x relationship for x < 100 MYA can be described well by a linear fit, with R2 values in the range 0.81-0.85. The linear fit (black line), equation and R2 values are shown for RGCs. MYA, million years ago.

We used a standardized computational pipeline to normalize, correct batch effects, reduce dimensionality, and cluster the data from each species separately(Stuart et al., 2019) (**Methods**). Biological replicates within each collection exhibited a high degree of concordance (**Extended Data Figs. 3-6)**. The numbers of cells in each class for each species are summarized in **Supplementary Table 1**.

**Fig. 3.**
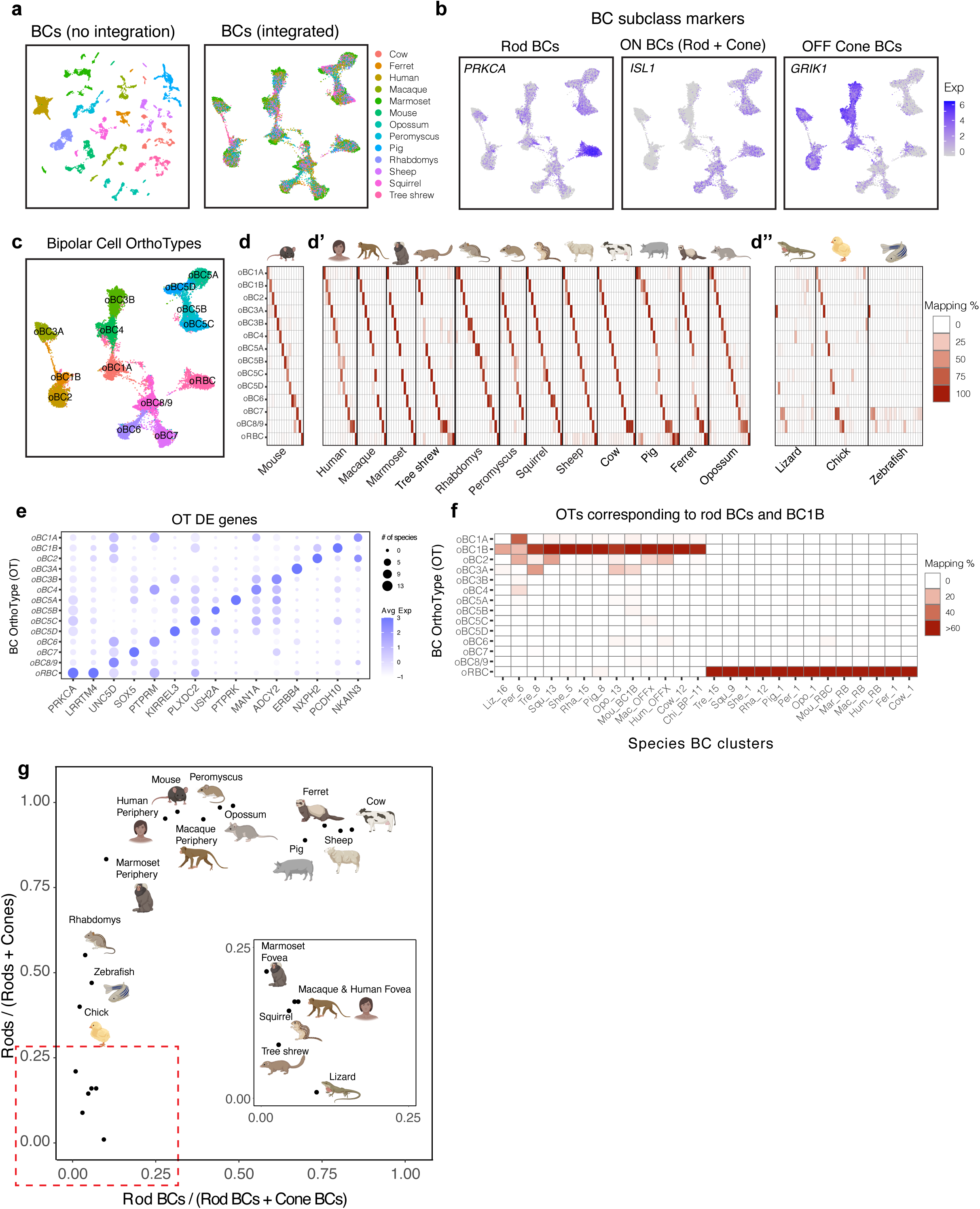
Multispecies integration of bipolar cells. a. UMAP of mammalian BCs computed with the raw (*left*) and integrated (*right*) gene expression matrices. Cells are colored by species of origin. b. Feature plots showing expression within the integrated space of a rod BC marker PRKCA (*left*), an ON BC marker *ISL1* (middle) and an OFF BC marker *GRIK1* (right). c. Same as the right panel of a, but with cells colored based on OrthoType (OT) identity as documented in **Extended Data Fig. 9a**. d. Confusion matrix showing specific mapping between mouse BC types and mammalian BC OTs. Each element represents the percentage of cells from a mouse BC type (column) that maps to a mammalian BC OT (row; see scale on the right of panel **d’’**). Each column sums to 100%. See **Extended Data Fig. 9a** for a higher magnification view. d’. Confusion matrix showing specific mapping between species-specific BC clusters (**Extended Data Figs. 3-6**) and mammalian BC OTs. Representation as in panel **d.** Columns are grouped by mammalian species demarcated by vertical black lines. d’’. Confusion matrices showing the mapping of BC clusters (columns) in lizard, chick and zebrafish to the mammalian BC OTs. Mapping that includes non-mammalian OTs is shown in **Extended Data Fig. 9e**. e. Dotplot showing differentially expressed genes (columns) within each BC OT (rows). The size of the dot represents the number of mammalian species (out of 13) that express the gene in at least 30% of cells mapping to the correspond OT, while the color represents normalized expression level. f. Confusion matrix showing the species BC clusters (columns) that map specifically to the OTs oRBC and oBC1B. BC types are named based on their species of origin and within-species BC cluster ID (**Extended Data Figs. 3-6**). For example, Peromyscus BC cluster 1 is called “Per_1”. g. Scatter plot showing the relative frequencies of rods among PRs (y-axis) vs. rod BCs among BCs (x-axis) for all jawed vertebrate species in this study (points, icons). For primates, data from fovea and periphery are plotted as separate points. The region demarcated by the dashed red box near the origin is expanded in the inset. RBC proportions were calculated from the atlases (**Extended Data Fig. 3-6**), while rod proportions were obtained from literature (see **Supplementary Note 1**).

## Molecular conservation of neuronal classes

We analyzed expression of class markers that have been validated in mice and primates – genes co-expressed within a retinal cell class with mutually exclusive expression across classes (Cowan *et al*., 2020; Macosko *et al*., 2015; Peng *et al*., 2019; Shekhar *et al*., 2016; Tran *et al*., 2019; Yan *et al*., 2020a; Yan *et al*., 2020b). Many showed similar expression patterns in other vertebrates (**Fig. 2a**). Using these markers, we assigned cells within each species to one of the six classes. We then assessed the interspecies similarity of classes by comparing “pseudobulk” transcriptomic profiles based on shared orthologous genes (**Methods**). A cross-correlation analysis among the 16 jawed vertebrates showed that transcriptomic similarity was driven by cell class identity rather than species identity – for example BCs of a given species are more closely related to BCs of other species than they are to other classes from the same species (**Fig. 2b,c and Extended Data Fig. 7a, b**). Qualitatively similar results were obtained when lamprey, a jawless vertebrate, was included, although the signal was attenuated as fewer orthologous genes were available (**Extended Data Fig. 7c-d**). Thus, class identity dominates species identity within a retinal cell’s transcriptional profile.

We next assessed evolutionary trends by comparing gene expression similarity and evolutionary distance among all pairs of species for each cell class. For all classes, interspecies transcriptomic correlation decreased with evolutionary distance (**Fig. 1b, 2e**). Among non-marsupial mammals, which separated less than 100 million years ago, the correlation decreased linearly with evolutionary distance (R^2^ = 0.827± 0.016), consistent with the predictions of neutral evolution (Chen *et al*., 2019). However, this decrease plateaued in comparisons with species that diverged >150 million years ago, suggestive of stabilizing selection (Chen *et al*., 2019). Interspecies distances within gene expression dendrograms for each cell class also reflected phylogeny (**Extended Data Fig. 7e**). Thus, major transcriptomic features of the six cell classes are conserved across vertebrates, consonant with their conserved morphology and connectivity. Moreover, cross-species transcriptomic variation is similar across cell classes, and is shaped by the species’ evolutionary history.

We found that conserved cell class genes included numerous known lineage-determining transcription factors (TFs) such as *POU4F1* (RGC), *VSX2* (BC and MG), *OTX2* (PR and BC), *TFAP2A-C* (AC), *ONECUT1/2* (HC), and *CRX* (PR) (Petridou and Godinho, 2022) (**Fig. 2a**). This suggests that the genetic mechanisms underlying neurogenesis and fate specification of cell classes are evolutionarily ancient.

## Molecular conservation of neuronal subclasses

Classically, three of the retinal cell classes have been subdivided into subclasses: PRs include rods, specialized for low-light vision, and cones, which mediate chromatic vision. Nearly all ACs use either γ-Aminobutyric acid (GABA) or glycine as their neurotransmitter, and transmitter choice is highly correlated with key morphological features. BCs can be subdivided into those that depolarize and hyperpolarize to illumination – ON and OFF types, respectively (Masland, 2012). Within PRs, ACs and BCs, cells from different species segregated based on subclass identity, and expressed orthologs of gene markers that have been well-characterized in mice **(Fig. 2d,d’** and **Extended Data Fig. 8a-c)**. Thus, the evolutionary conservation of cell classes extends to subclasses.

Several transcription factors are expressed selectively in mouse retinal subclasses, including *NRL* and *NR2E3* in rods, *THRB* and *LHX4* in cones, *MEIS2* in GABAergic ACs, *TCF4* in glycinergic ACs, *FEZF2* and *LHX3* in OFF BCs, and *ISL1* and *ST18* in ON BCs (Petridou and Godinho, 2022). Some, including *NRL*, *NR2E3*, *THRB* and *ISL1*, have been implicated in the differentiation of the subclass that expresses them. The subclass-specific expressions of these TFs were broadly conserved across species **(Extended Data Fig. 8d)**, suggesting that programs specifying subclasses, like those specifying classes, are evolutionarily ancient **(Extended Data Fig. 8e-f)**.

## Tight conservation of mammalian bipolar cell types

We next considered the conservation of neuronal types within classes. We began by analyzing the evolutionary variation among mammalian BC types, which have been comprehensively classified in mice (Shekhar *et al*., 2016), macaques (Peng *et al*., 2019) and humans (Yan *et al*., 2020b). In mice, for example, there are 15 BC types: 6 OFF and 9 ON BC types; one of the ON BC types receives input predominantly from rods (RBCs) and all others predominantly from cones (Shekhar *et al*., 2016).

Initial clustering of mammalian BCs generated groups that were defined by species (**Fig. 3a**). The datasets were therefore reanalyzed using an integration method that minimizes species-specific signals, thereby emphasizing other transcriptomic relationships (**Methods**) (Stuart *et al*., 2019). This analysis intermixed the species, while retaining structure that separates ON cone, OFF cone, and ON RBCs from each other (**Fig. 3b**).

The integrated data revealed 14 groups of cells based on shared transcriptomic signatures (**Fig. 3c**). Even though species-specific cluster labels were not an input to the analysis, mouse BC types mapped to the integrated groups in a 1:1 fashion, with the sole exception of two closely related and sparsely represented types (BC8/9) that mapped to the same group (**Fig. 3d and Extended Data Fig. 9a**). We call these groups neuronal OrthoTypes (OTs) although, as in the case of BC8/9, they may sometimes contain small sets of related types. We named the BC OTs by the mouse types; thus, the OT containing mouse BC1A is called oBC1A, and so on. Each BC OT was represented in nearly all mammals (**Extended Data Fig. 9b**), and 91% of mammalian BC clusters (172/190) mapped specifically to a single OT (**Fig. 3d’**). We identified differentially expressed genes that distinguished the BC OTs (**Fig. 3e**).

The “mammalian” OTs remained robust when mammalian, chick, lizard and zebrafish BCs were integrated together. Although 32% fewer orthologous genes were available to guide the analysis, many BC clusters in chick, several in lizard, and a few in zebrafish mapped to these mammalian OTs (**Fig. 3d’’**). However, two additional “non-mammalian” OTs emerged, comprising OFF BCs and ON BCs from the non-mammals (**Extended Data Fig. 9c-e)**. Attempts to find additional substructure in these non-mammalian BC OTs were unsuccessful, likely because chick, lizard and zebrafish are nearly as evolutionarily distant from each other as they are from mammals. Nonetheless, the fact that several chick and lizard BC clusters map to the mammalian OTs suggests that some type-specific BC identities have been conserved for >300 million years.

To illustrate the utility of the integration, we highlight two BC OTs – oRBC and oBC1B (**Fig. 3f**). RBCs receive most of their input from rods, as their name implies, and they connect with specific AC types rather than directly with RGCs (Grimes et al., 2018). oRBC contained RBCs from all mammals (**Fig. 3f**). Mammalian RBCs were distinguished by the high expression of the genes *PRKCA* and *LRRTM4* (**Fig. 3e**) both of which are RBC-specific in mice (Shekhar *et al*., 2016). RBCs also exhibit species-specific gene expression (**Extended Data Fig. 9f**). RBCs have been described in chicks (Yamagata *et al*., 2021) and zebrafish (Kolsch *et al*., 2021), but these types did not map to oRBC.

Having identified rods as well as RBCs, we could ask whether their abundances (rods as a fraction of all PRs and RBCs as a fraction of all BCs) were related. Some retinas (e.g. squirrel and lizard) and retinal regions (primate fovea) are highly cone-dominant while others (e.g., mouse, ungulates and primate peripheral retina) are rod-dominated. RBC abundance is thought to track rod abundance and quantitative analysis confirmed this trend but showed an intriguing division into two groups: in one group, rods varied over a wide range (1-55% of all PRs) and RBCs comprised <10% of all BCs, whereas in a second group, rods comprised >75% of PRs and RBCs varied over a wide range (15-85% of BCs) (**Fig. 3g**).

The second OT represents a non-canonical OFF BC recently described in mice, and named BC1B (Shekhar *et al*., 2016) or GluMI (Della Santina et al., 2016). The name BC1B reflects its transcriptional similarity to BC1A. However, unlike canonical BCs, BC1B retracts its dendrite during early postnatal life and therefore has no direct connection with mature PRs (Shekhar *et al*., 2016). Likely because it lacks this canonical feature, no BC1B equivalent has yet been identified in other species. However, 10 of the 13 mammals profiled here, as well as chicks and lizards, contained a BC cluster that mapped exclusively to oBC1B **(Fig. 3f**), while two mammals (Peromyscus and ferret) contained a cluster that mapped to both oBC1A and oBC1B. Thus, transcriptomics allowed identification of a potentially conserved cell type that would have been difficult to identify by conventional morphological methods; its type-specific markers can now be used to seek morphological and physiological validation.

## Retinal ganglion cell OrthoTypes

We next performed OT analysis on RGCs, the retina’s sole output neurons. We identified 21 RGC OTs in mammals, and found differentially expressed (DE) genes that distinguished them (**Fig. 4a-c** and **Extended Data Fig. 10a)**. 81% of mammalian RGC clusters (329/408) mapped predominantly to a single OT (**Fig. 4d)**. In species that contain more RGC types than OTs, transcriptomically similar RGC clusters mapped to the same OT. As was the case for BCs, RGC OTs remained stable when lizard, chick and zebrafish were included in the integration (**Fig. 4d’)**, but were supplemented by an additional OT dominated by non-mammalian species (**Extended Data Fig. 10b-d**).

**Fig. 4.**
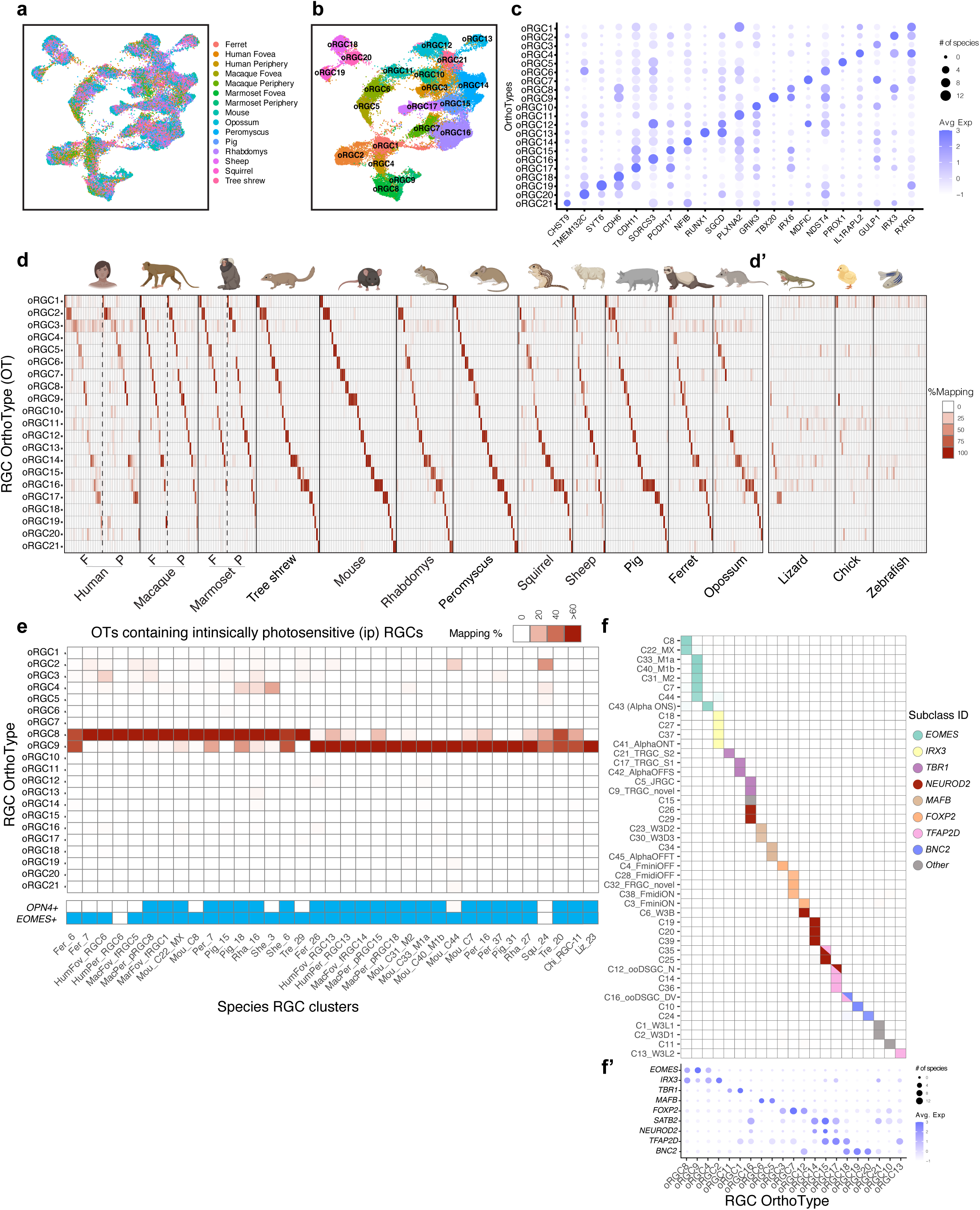
Multispecies integration of retinal ganglion cells. a. Integrated UMAP of RGCs from 12 mammals (Cow was excluded due to paucity of RGC data.). Cells are labeled by species of origin. For primates, cells from fovea and periphery are plotted separately. b. Same as **a**, with RGCs labeled by OT. c. Dotplot showing differentially expressed genes (columns) within each RGC OT (rows). Representation as in **Fig. 3e**. d. Confusion matrices showing that species-specific RGC clusters (**Extended Data Fig. 3-6**) map to mammalian RGC OTs in a specific fashion. Representation as in **Fig. 3d’** except that clusters from Fovea (F) and Periphery (P) are mapped separately for primates. d’. Confusion matrices showing the mapping of RGC clusters (columns) in lizard, chick and zebrafish to the 21 mammalian RGC OTs. Mapping to the single non-mammalian RGC OT is shown in **Extended Data Fig. 10d**. e. Confusion matrix showing the species-specific RGC clusters (columns) that map to the oRGC8 and 9, corresponding to ipRGCs. Representation as in **Fig. 3f**. Annotation bar (bottom) highlights species-specific RGC clusters that express *OPN4* and *EOMES*, a transcription factor expressed selectively by ipRGCs (Rheaume *et al*., 2018; Tran *et al*., 2019). f. Confusion matrix showing that mouse RGC types (rows; naming as in ref. 19) belonging to TF-based subsets (Shekhar, *et al*., 2022) (colors) map to the same OTs (columns). f’. Dotplot showing specific expression patterns of subclass-specific TFs (Shekhar, *et al*., 2022) in OTs. Representation as in **Fig. 3e**.

Because mapping between types and OTs was more complex for RGCs than for BCs (see below), we tested the integration by seeking orthologs of an evolutionarily ancient set of RGC types called intrinsically photosensitive (ip) RGCs. ipRGCs contain the photopigment melanopsin (*OPN4*), which allows them to generate visually evoked signals without input from PRs (Hattar et al., 2002). They mediate crucial non-image forming visual functions such as circadian entrainment and the pupillary light reflex. ipRGCs have been detected in the retinas of diverse vertebrate orders, including several of the species profiled here, generally on the basis of *OPN4* expression (Do, 2019). ipRGCs also expresses the transcription factor *EOMES/TBR2*, although some *EOMES*-expressing RGCs have not yet been functionally validated as ipRGCs. RGCs in two OTs, oRGC8 and oRGC9, expressed *OPN4* (**Fig. 4e**). oRGC9 contained 5 mouse RGC types, three of which were the ipRGC types M1a, M1b and M2, which express the highest levels of melanopsin. oRGC8 contained the paralogous types, MX and C8. Overall, of 35 clusters from 11 species in these two oRGCs, 25 expressed *OPN4* and 33 expressed *EOMES*. *OPN4*-expressing RGC types from chick and lizard also mapped to these OTs. Thus, cross-species integration captures an RGC group with a conserved physiological property.

We showed recently that 45 molecularly-defined mouse RGC types, many of which map to physiologically and morphologically-defined mouse RGC types (Goetz et al., 2021), can be grouped into subsets defined by selectively expressed TFs (Shekhar et al., 2022; Tran *et al*., 2019; Whitney et al., 2023). Some of these TFs (e.g. *EOMES*, *TBR1*, and *NEUROD2*) have been implicated in RGC development (Cherry et al., 2011; Kiyama et al., 2019; Liu et al., 2018; Mao et al., 2014). The mapping of mouse RGC types to RGC OTs mirrored these subsets (**Fig. 4f**), and subset-defining TF expression patterns were recovered in a large proportion of species (**Fig. 4f’**). These results suggest that, as noted above for PR, BC and AC subclasses, it may be possible to classify RGCs into evolutionarily conserved subclasses.

Although RGCs and BC OTs were represented in all mammals, the number of neuronal types within a species varied over a greater range for RGCs than for BCs (28 ± 11 [mean ± SD] vs 14 ± 2 types) (**Extended Data Fig. 3-6**). While a BC OT was associated with 1.37 ± 0.5 BC types within a species, an RGC OT was associated with 2.55 ± 2 RGC types. Thus, type variation is greater for RGCs than BCs. Although we did not conduct formal OT analysis of PR and HC types, they are highly conserved (Masland, 2012). There are only 3 PR types in most mammals - rods, S-cones (short wavelength sensitive) and M/L-cones (medium and long wavelength sensitive), with trichromats having separate M- and L-types that differ only in the opsin they express (Peng *et al*., 2019). Most mammals have 2 HC types, one with and one without an axon; mice have only a single type, which is axon-bearing. Although ACs are poorly annotated and cannot be integrated across species at this time, it is clear that divergence of RGC types is greater than the other three retinal neuronal classes.

## Orthologs of midget and parasol RGCs

In most species studied to date, no RGC type comprises more than ∼10% of all RGCs. In striking contrast, the retina of many primates, including humans, is dominated by two closely-related RGC types, ON and OFF midget RGCs, named for their diminutive somata and dendritic trees (Polyak, 1941). Together they account for >80% of all RGCs in macaque and human, with similar abundance in fovea and periphery (Peng *et al*., 2019; Yan *et al*., 2020b). However, despite their importance for vision, no non-primate orthologs of midget RGCs have been found, and our own prior comparison of mouse and macaque primate RGCs failed to find any correspondence (Peng *et al*., 2019). Similarly, attempts to find orthologs of the next most abundant primate RGC types, ON and OFF parasol RGCs (5-10% of all RGCs) have remained inconclusive (Berson, 2008).

We used OTs to revisit this issue. Each of the four abundant primate types mapped to a distinct OT (oRGC1, 2, 4 and 5), and each of these OTs contained the corresponding cell type from both fovea and periphery of human, macaque, and marmoset (**Fig. 5a** and **Extended Data Fig. 11a**). Remarkably, the mouse RGC types mapping to these OTs included a set of four related types called α-RGCs (Krieger et al., 2017); they accounted for 3 of the 5 mouse types in the midget- and OFF parasol-containing OTs. The resemblance of parasol RGCs to α-RGCs has been noted previously (Crook et al., 2008; Peng *et al*., 2019), but the correspondence was particularly unexpected for midget RGCs, because α-RGCs are low abundance (<2% per type) and among the largest mouse RGCs. Nonetheless, several lines of evidence support the significance of the orthology between primate midgets and parasols, and the mouse α-RGC types.

**Fig. 5.**
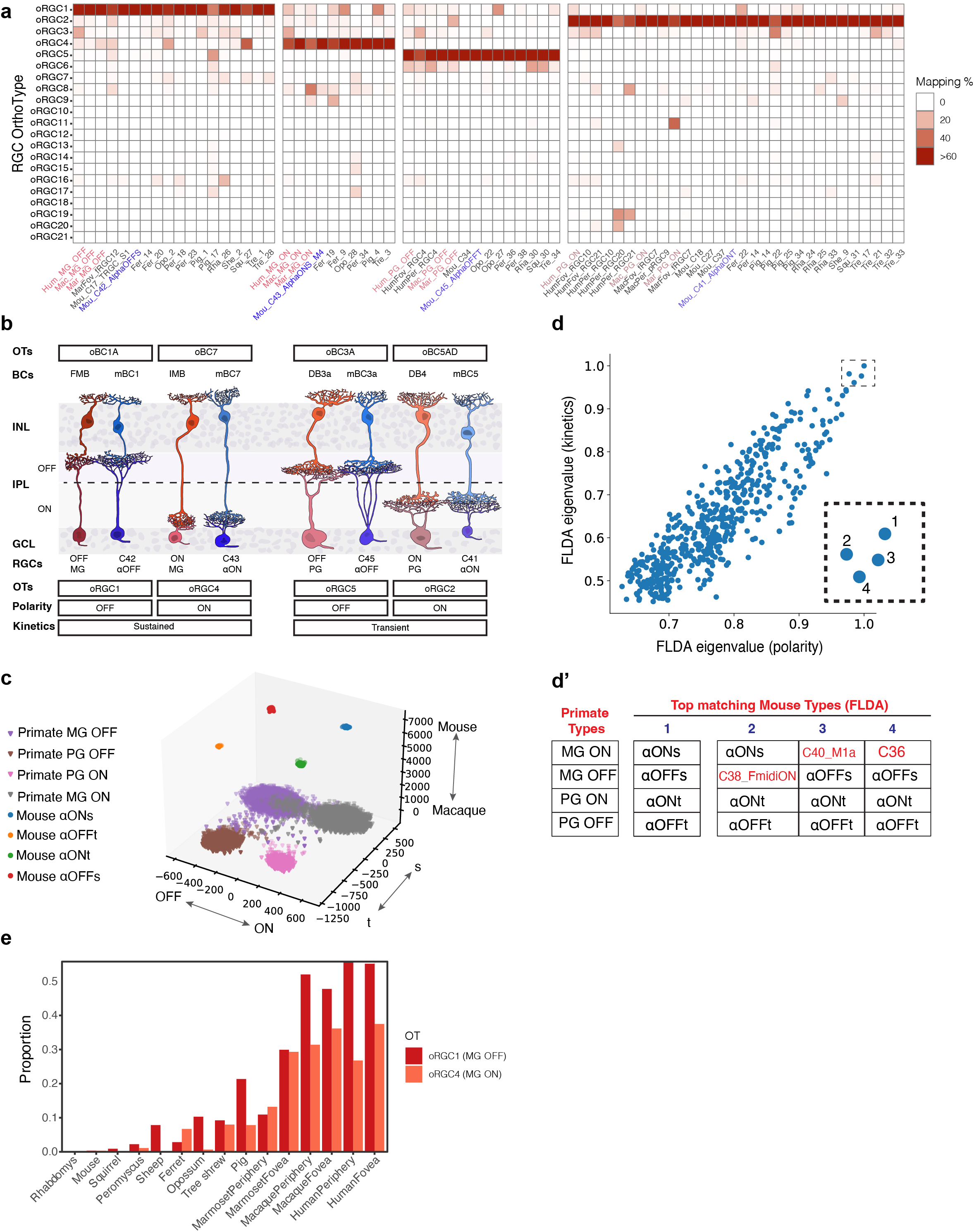
Mammalian orthologs of midget and parasol RGCs. a. Confusion matrix showing RGC clusters from each species (columns) that map specifically to oRGC1, oRGC4, oRGC5 and oRGC2, which respectively contain OFF and ON midget RGCs (MG), and OFF and ON parasol RGCs (PG). Representation as in **Fig. 3f**. Columns corresponding to primate midget and parasol types are shown in red, and mouse α-RGC types are shown in blue. b. Schematic delineating morphological and physiological similarities between primate/midget RGCs and their α-RGC orthologs. OrthoTypes of each pair as well as the orthology among BC types that innervate them is also shown. Morphologies of neuronal types were created based on published data (**Supplementary Note 2**). Within each pair, the left column corresponds to primate types and the right column corresponds to mouse types. c. FLDA projection of the scRNA-seq data for primate midget and parasol, and mouse α-RGC types onto the corresponding three-dimensional space with axes representing species, polarity and kinetics. d. Scatter plot of the FLDA eigenvalues for the kinetics (y-axis) vs. polarity (x-axis), measuring the magnitude of the variance corresponding to these attributes captured in the projection. Inset highlights the top four matches. d’. Mouse RGC types present within the top four matching combinations with primate midget and parasol RGCs. e. Relative proportion of OFF and ON midget RGC orthologs in mammalian species based on the frequencies of cells in oRGC1 and oRGC4.

First, the four α-RGC types can be distinguished based on response polarity (ON vs. OFF) and response kinetics (sustained [s] vs. transient [t]): α-ONs, α-OFFs, α-ONt, and α-OFFt(Krieger *et al*., 2017). Mouse α-ONs and α-OFFs mapped to ON and OFF midgets respectively, and mouse α-ONt and α-OFFt mapped to ON and OFF parasols respectively. Second, midgets and parasols exhibit sustained and transient light responses, respectively, which match the kinetics of their mouse orthologs (de Monasterio, 1978; Krieger *et al*., 2017). Third, dendrites of matched types laminate to similar sublaminae in the inner plexiform layer, with the parasol and α-transient types nearer the center of the layer than the midget and α-sustained types (Nassi and Callaway, 2009). Fourth, morphological studies have identified the BC types that innervate midgets, parasols and α-RGCs (Tsukamoto and Omi, 2015; 2016; Yu et al., 2018). In each case, the primate BC type that provides the majority of excitatory input to the midget or parasol RGC type is a member of the same BC OT as a mouse BC type that provides substantial input to the corresponding α-RGC type. Thus, although none of these metadata were provided explicitly, the integration matched types correctly based on their polarity, response kinetics, dendritic lamination and inputs (**Fig. 5b**). In addition, primate midget/parasol and mouse α-RGCs exhibit similar response properties: midget RGCs and sustained α-RGCs, primarily report on contrast and are minimally feature-selective, whereas parasol RGCs and transient α-RGCs, are motion-sensitive (Manookin et al., 2018; Wang et al., 2021).

We used the parallels in kinetics and polarity to assess the strength of the midget/parasol – α-RGC correspondence with an independent statistical approach, Factorized Linear Discriminant Analysis (FLDA) (Qiao and Meister, 2020) (**Extended Data Fig. 11b**). Given single-cell transcriptomic data from cells that carry multiple categorical attributes, FLDA attempts to factorize the gene expression data into a low dimensional representation in which each axis captures the variation along one attribute while minimally co-varying with other attributes. We applied FLDA to project primate midgets and parasols, and mouse α-RGCs onto a 3-D space whose three axes represent species (mouse-primate), kinetics (sustained-transient), and polarity (ON-OFF). FLDA generated a projection in which the relative arrangement of the four primate and the four mouse cell types was consistent with their attributes (**Fig. 5c and Extended Data Fig. 11c**). We then asked whether α-RGCs were a better transcriptomic match to midgets and parasols than other mouse RGC types carrying similar attributes. For this purpose, we identified a set of 20 mouse RGC types for which polarity (ON/OFF) and kinetics (sustained/transient) are known (**Supplementary Table 2**). We matched all possible 432 combinations of 4 drawn from this set with the midgets and parasols, calculated the FLDA projections, and ranked them based on the magnitude of the variance captured by FLDA along the polarity and kinetics axes (**Fig. 5d**). The best match comprised all four α-RGC types, and the next three matches contained three α-RGC types plus one other type (**Fig. 5d’**). This result provides strong support for the orthology of primate midget and parasol RGCs with mouse α-RGCs, suggesting that the former are not primate innovations, as they have been considered to be. Moreover, the presence of midget and parasol orthologs in all the mammals studied here (**Fig. 5e** and **Extended Data Fig. 11d**) suggests that they are likely to have evolved from antecedent types present in the mammalian common ancestor.

For midget RGCs, we suggest a relationship between their dramatic expansion in the primate lineage (**Fig. 5e**) and the evolution of visual processing. In primates, the principal retinorecipient region in primates is the dorsolateral geniculate nucleus (dLGN), whereas in mice it is the superior colliculus (SC) (Seabrook et al., 2017). Midget RGCs project almost exclusively to the dLGN (Dacey et al., 2003). In mice, anterograde (Martersteck *et al*., 2017) and retrograde (Johnson et al., 2021; Rosón et al., 2019) tracing studies suggest that α-RGCs are overrepresented among those RGCs that project to the dLGN (2-4 fold in Rosón et al., 2019). The dLGN provides the dominant visual input to the primary visual cortex, whereas SC projects in large part to areas that control reflexive motor responses including eye movements (Ito and Feldheim, 2018). In primates, complex visual processing occurs largely at the cortical level, and may be best served by the relatively unprocessed, high-acuity rendering of the visual world that midget RGCs provide. The modest loss in response time in this system is presumably compensated by the greater flexibility in response type. As the cortex plays a key role in primate vision, midget-like RGCs already present in the mammalian ancestor may have increased in number and decreased in size in primates to facilitate this flexibility.

## Conclusions

We integrated single-cell transcriptomic cell atlases of the retina from 17 vertebrate species and used them to assess the extent to which cell classes, subclasses and types have been conserved through vertebrate evolution. Our main results and the conclusions we draw from them are as follows: **First**, retinal cell classes and subclasses are highly conserved at the molecular level through evolution, mirroring their structural and functional conservation. The pattern of gene expression variation in classes exhibits signatures of neutral drift within mammals and of stabilizing selection in the evolution from non-mammalian vertebrates to mammals. **Second**, although greater cross-species variation exists at the level of cell types, numerous conserved types can be detected using an analytical framework that identifies transcriptomic groups, which we call OrthoTypes. **Third**, evolutionary divergence among types is more pronounced for RGCs than other retinal classes, suggesting that the outer retina is built from a conserved parts list while natural selection acts more strongly on diversifying those neuronal types that transmit information from the retina to the rest of the brain. **Fourth**, conserved transcription factors at all three levels (class, subclass, type) suggest that developmental programs for the specification of retinal neurons may have an ancient origin. **Fifth**, midget and parasol RGCs, which together comprise >90% of human RGCs, have orthologs in other mammalian species, suggesting that these primate cell types are derived from the expansion and modification of types present >300 million years ago in the retina of the last common ancestor of mammals. In mice, the orthologs are a numerically minor set of types called α-RGCs. The dramatic (>20-fold) difference in abundance of midget orthologs between mice and humans correlates with the greater prominence of visual processing in the primate cortex. Finally, knowing the orthologs of midget and parasol RGCs in several accessible models will aid efforts to slow their degeneration in blinding diseases such as glaucoma.

## Acknowledgements

This work was supported by the NIH (K99EY033457, R00EY028625, R21EY028633, U01MH105960 and T32GM007103), the Chan-Zuckerberg Initiative (CZF-2019-002459), the Glaucoma Research Foundation (K.S.), startup funds from the UC Berkeley (K.S.), an award from Research to Prevent Blindness and a Klingenstein-Simons Fellowship Award (Y.R.P.), a Wellcome Trust Investigator Award (210684/Z/18/Z) (R.J.L.), an ARCS Foundation Scholarship and a Society for Developmental Biology Emerging Models grant (A.M.R.), and grants from Children’s Glaucoma Foundation and NSF (1827647) to James Lauderdale and Douglas B. Menke. We thank James D. Lauderdale and Douglas B. Menke (University of Georgia) for supervision of A.M.R.; Mallory Laboulaye and Rebecca Schaffer for assistance; Joseph Wekselblatt for tree-shrew tissue.; Guoping Feng for marmoset tissue; Hopi Hoekstra for peromyscus tissue; and Steve Van Hooser for ferret tissue. Icons for species in the figures were obtained from BioRender.com.

## Author Contributions

J.R.S and K.S. conceived the study and supervised the project. J.H., W.Y., K.S., and A.Ku. performed the computational analysis. A.M. performed the sc/snRNA-seq and histology experiments with contributions from A.Ka., Y.K., and Y.R.P.. A.M.R, R.R., H.B., R.J.L, W.L, and J.T.T. provided tissue. M.Q. performed the FLDA analysis, and was supervised by M.M. R.J.L. and R.R. provided an annotated Rhabdomys genome. J.H., A.M., J.R.S. and K.S. wrote the paper with input from other authors.

## Corresponding authors

Correspondence to Joshua R. Sanes and Karthik Shekhar.

## Ethics Declarations

<u>Competing interests:</u> The authors declare no competing interests.

## Supplementary Information

**Supplementary Note 1.** Literature sources used to derive the proportion of rods among PRs.

**Supplementary Note 2.** Literature sources used for mouse and primate cell type morphologies shown in **Fig. 5b**.

**Supplementary Note 3.** Mathematical background for Factorized Linear Discriminant Analysis (FLDA).

**Supplementary Table 1**. Numbers of cells/nuclei per retinal class per species that passed quality metrics. Note that the proportions do not represent endogenous values, but reflect the results of antibody-based enrichment, which was directed to recover BCs and RGCs in many samples.

**Supplementary Table 2**. List of mouse RGC types with known polarity and kinetics tested for matching with primate ON/OFF midget and ON/OFF parasol RGC types using Factorized Linear Discriminant Analysis (FLDA).

**Supplementary Table 3.** List of reference transcriptomes used to align scRNA-seq and snRNA-seq data from species analyzed in this study.

## Materials and Methods

### Ethical Compliance

Human eyes were obtained post-mortem at a median of 6 hours from death either from Massachusetts General Hospital (MGH) via the Rapid Autopsy Program or from The Lion’s Eye Bank in Murray, Utah. Acquisition and use of post-mortem human tissue samples were approved by either the Institutional Review Board of the University of Utah, or the Human Study Subject Committees of Harvard Medical School, and they were in compliance with the National Human Genome Research Institute (NHGRI) policies. Informed consent was obtained from participants if they were enrolled antemortem, or their legal guardians if post-mortem. All donors were confirmed to have no history or clinical evidence of ocular disease or intraocular surgery. Pig, cow and sheep eyes were obtained, on average, 1 hour from death from an abattoir located in West Groton, Massachusetts. Other animal eyes were obtained from animal colonies maintained at Brandeis University (ferret), California Institute of Technology (tree shrew), Harvard University (ferret), MIT (marmoset), NIH (squirrel), University of Manchester, UK (Rhabdomys), University of Georgia (lizard), and University of California, Los Angeles (lamprey, opossum). Animal experiments conducted in the United States were approved by the Institutional Animal Care and Use Committees (IACUC) in each location. Rhabdomys tissue was collected in accordance with the Animals, Scientific Procedures Act of 1986 (United Kingdom) and approved by the University of Manchester ethical review committee.

### Single nucleus RNA sequencing

#### Nuclei-isolation and sorting

For isolation of nuclei, frozen retinal tissues were homogenized in a dounce homogenizer in 1ml lysis buffer consisting of 0.1% NP-40 in a solution containing 10 mM Tris, 1mM CaCl_2_, 8mM MgCl_2_, 15mM NaCl, 0.1U/µl RNAse inhibitor (Promega RNasin Ribonuclease Inhibitor N2615), and 0.02U/µl DNAse (D4527, Sigma Aldrich). The homogenized tissue was passed through a 40-µm cell strainer. The filtered nuclei were pelleted at 500 rcf for 5 min, resuspended in staining buffer (Tween 0.02%, and 2% BSA in the tris base buffer) and stained with anti-NEUN (1:300, Sigma #FCMAB317PE or #MAB377A5) and anti-CHX10 (1:600, Santa Cruz Biotechnology #sc-365519 AF647) for 12min at 4°C.

Following staining, nuclei were centrifuged, resuspended in sorting buffer (2% BSA in the Tris base buffer), and counterstained with DAPI (1:1000). The NEUN+ and CHX10+ nuclei were sorted into separate tubes (**Extended Data Fig. 2a-c**), pelleted again at 500 rcf for 5 min, resuspended in 0.04% non-acetylated BSA/PBS solution, and adjusted to a concentration of 1000 nuclei/µL. The integrity of the nuclear membrane and presence of non-nuclear material were assessed under a brightfield microscope (**Extended Data Fig. 2d,e**) before loading into a 10X Chromium Single Cell Chip (10X Genomics, Pleasanton, CA) with a targeted recovery of 8000 nuclei per channel.

#### Library preparation

Single nuclei libraries were generated with either Chromium 3’ V3, or V3.1 platform (10X Genomics, Pleasanton, CA) following the manufacturer’s protocol. Briefly, single nuclei were partitioned into Gel-beads-in-EMulsion (GEMs) where nuclear lysis and barcoded reverse transcription of RNA would take place to yield cDNA; this was followed by amplification, enzymatic fragmentation and 5’ adaptor and sample index attachment to yield the final libraries. Libraries were sequenced on an Illumina NovaSeq at the Bauer Core Facility at Harvard University. Sequencing data were demultiplexed and aligned using Cell Ranger software (version 4.0.0, 10X Genomics, Pleasanton, CA).

### Histology

Whole eyes were fixed in 4% paraformaldehyde (in PBS) for 1-2 hour and then transferred to PBS. Either whole retinas or 8mm punches of central retina were dissected out and sunk in 30% sucrose in PBS overnight at 4°C, before being embedded in tissue freezing medium and sectioned coronally at 20 µm in a cryostat. Sections were mounted onto coated slides. Slides were incubated for 1 hour with 5% donkey serum (with 0.1% TritonX) at room temperature, then overnight with primary antibodies (1:500 RBPMS, PhosphoSolutions #1832-RBPMS; 1:400 CHX10, Novus Biologicals #NBP1-84476; 1:50 AP2A, DSHB #3B5) at 4°C, and finally for 2 hours with secondary antibodies in PBS at room temperature. Images were acquired on Zeiss LSM 900 confocal microscopes with 405, 488, 568, and 647 nm lasers, and processed using Zeiss ZEN software suites.

### Preprocessing of transcriptomic data

We used cellranger (v7.0, 10X Genomics) to align the sc- and snRNA-seq datasets, following manufacturer’s instructions. For each species, sequencing reads were demultiplexed into distinct samples and the .fastq.gz files corresponding to each sample were aligned to reference transcriptomes to obtain binary alignment map (.bam) files. The reference transcriptomes used are listed in **Supplementary Table 3**. To include both exonic and intronic reads in the quantification of gene expression for each sample, regardless of cellular or nuclear origin, we applied velocyto(La Manno et al., 2018) to the corresponding .bam files. This generated two separate gene expression matrices (GEMs; genes x cells) for each sample, corresponding to “spliced” and “unspliced” reads. The two GEMs were summed element by element to obtain “total” GEM for each sample. For each species, GEMs from different samples were combined (column-wise concatenated) to yield a species’ GEM.

### Computational Analysis

Analysis of the GEMs was performed in R. While several packages were used for statistical calculations and data visualizations, our workflow was based on the Seurat package for single-cell analysis developed and maintained by the Satija Lab (Hao et al., 2021; Stuart *et al*., 2019) (https://satijalab.org/seurat/). We describe the analysis steps here at a high-level. We have also made available the analysis scripts and processed datasets, including annotated Seurat objects, on our Github page (www.github.com/shekharlab).

#### Segregation of major retinal cell classes

Data from each species were separately analyzed through a clustering procedure to identify high-quality cells, and segregate the major cell classes (PR, BC, HC, AC, RGC, MG). Briefly, GEMs from different replicates were combined, and transcript counts in each cell was normalized to a total library size of 10,000 and log-transformed (X ←log(X+1)). We identified top 2000 highly variable genes (HVGs), and applied principal component analysis (PCA) to factorize the submatrix corresponding to these HVGs. Using the subspace corresponding to the top 20 principal components (PCs), we built a k-nearest neighbor graph on the data, and then clustered with a resolution parameter of 0.5 using Seurat’s FindClusters function. The same PCs were used to embed the cells onto a 2D visualization using the Uniform Manifold Approximation (Becht et al., 2019). The 2D embeddings were solely used to visualize clustering structure and gene expression patterns *post hoc*.

Each cluster was assigned to one of the six major retinal cell classes based on expression of orthologs of canonical markers characterized in mice (Macosko *et al*., 2015): PRs (*Arr3*, *Rho*, *Crx*), HCs (*Calb1*, *Onecut1*, *Onecut2, Lhx1*), BCs (*Vsx1*, *Otx2*, *Grik1*), ACs (*Gad1*, *Gad2*, *Tfap2a*, *Tfap2b*, *Tfap2c*), RGCs (*Rbpms*, *Nefl*, *Nefm*, *Slc17a6*), and MG (*Glul*, *Apoe*, *Rlpb1*). Clusters that mapped to other cell types found at much lower frequency (e.g. endothelial cells, microglia) or which contained low quality cells were not considered further. The number of cells of each class in each species is indicated in **Supplementary Table 1**. We note that because many experiments were designed to enrich certain classes (RGCs or BCs), the relative frequencies do not reflect endogenous values.

#### Integration and clustering to to identify BC and RGC types

Given the focus on BC and RGC diversity in this work, we separated RGCs and BCs within each species, and clustered them independently using the following procedure. After subsetting the data by class (BC or RGC), cells with abnormally high ( > mean + 2*SD) or low (< mean - 2*SD) counts were removed. We also removed replicate batches that contained the class of interest at a frequency less than 50 cells. We split the cells by replicate ID and used Seurat’s integration pipeline to remove batch effects, reduce dimensionality and cluster the data in a shared low-dimensional integrated space. We selected 20-25 latent variables in the integrated space to identify clusters and generate 2D UMAP visualizations.

We initially deliberately overclustered the data using a resolution parameter of 1.1; clusters were then merged or pruned as follows: For each cluster, we calculated differentially expressed (DE) markers, and these markers were inspected to determine if clusters should be merged or removed. Some clusters were also removed if their top DE markers were widely expressed in several clusters, if they had lower RNA counts compared to other clusters, or if several of the top DE markers were canonical markers for contaminant cell classes. If more than 20% of cells were removed via pruning, the filtered data was subjected to another round of integration and clustering. Two or more clusters were merged if a differential expression test failed to find markers that sufficiently distinguished the clusters.

We applied these steps to define BC and RGC clusters for species initially reported in this paper: Peromyscus, Ferret, Opossum, Brown anolis lizard, Cow, Sheep, Pig, Thirteen-lined ground squirrel, Four-striped grass mouse, and Tree shrew. Individual clusters correspond to individual cell types, and in some cases, to small groups of closely related types. For the sake of consistency, we also applied the same procedure to RGC and BC data of species published elsewhere (Mouse (Shekhar *et al*., 2016; Tran *et al*., 2019), Macaque (Peng *et al*., 2019), Human (Yan *et al*., 2020b), Zebrafish (Kolsch *et al*., 2021), and Chick (Yamagata *et al*., 2021)). In all cases, our clusters were largely consistent with published annotations, and we therefore labeled these clusters based on their published labels.

#### Selection of shared orthologous genes

Orthologous genes were identified using orthology tables via Ensembl BioMart (https://useast.ensembl.org/info/data/biomart/index.html). Using mouse as a reference species, pairwise orthology tables were generated between mouse and every other species. These orthology tables contained information about the number of predicted orthologs for every mouse gene within each species. Mouse genes that had a 1:1 ortholog in every other species were retained as the set of orthologous features, with the exception of zebrafish. Due to a whole gene duplication, zebrafish has several paralogous pairs of genes (e.g. *rbpms2a* and *rbpms2b*) known as “ohnologs” (Howe et al., 2013). The prevalence of ohnologs results in a paucity of 1:1 orthologs. To address this issue, we collapsed each ohonolog pair by summing over their expression (e.g. *rbpms2a* and *rbpms2b* to *rbpms2*). If the ohnologs were the only orthologs of a gene, then the composite gene was regarded as the 1:1 ortholog for further analysis. Overall, we found 1905 1:1 orthologs among all 17 species, 4552 among the 16 jawed vertebrates (i.e., omitting lamprey) and 6693 among the 13 mammals. The number of shared orthologs decreased with evolutionary distance, and we found fewer orthologs shared between mammals and non-mammalian vertebrates than among mammals.

#### Analysis of cell classes and subclasses

For each species, we computed cell-averaged (or pseudobulk) gene expression vectors for the six major cell classes (PR, HC, BC, AC, RGC and MG). Spearman cross-correlation matrices were calculated for all mammals, the 16 jawed-vertebrates and all 17 vertebrates using the corresponding sets of orthologous genes. To analyze evolutionary trends within a class (**Fig. 2e**), we first calculated pairwise evolutionary distances in millions of years (MYA) using TimeTree (Kumar et al., 2017). These distances were also used to build the species dendrogram in **Fig. 1b**. We then plotted the Spearman correlation between every pair of species vs. evolutionary distance. We used locally estimated scatterplot smoothing (LOESS) to draw a trendline through the data using 2^nd^ degree polynomials. To fit data for evolutionary distances under 100 MYA, we used linear least squares regression. Dendrograms for the cell-averaged profiles within each class were constructed using the function hclust (package “stats”) using correlation distance, and then the cophenetic distances for each pair of species were computed using the function cophenetic (package “stats”).

For an alternative view on the cell classes, we subsampled each cell class to 200 per species, and then combined the GEMs. The resulting GEMs were integrated using Seurat using each species as a “batch”. Note that batch correction was not performed for samples within a species, nor was cell class information provided to the integration. The resulting integrated data was visualized on a UMAP (**Fig. 2d** and **Extended Data Fig. 8**). Dendrograms for the cell-averaged profiles were constructed using hclust (package “stats”), and then plotted in a circular representation using the circlize_dendrogram function (package “dendextend”) (**Extended Data Fig. 7a**).

#### Data integration and identification of OrthoTypes

We identified OrthoTypes (OTs) separately for BCs and RGCs. In each case, we followed the following steps: (i) Within each species, the corresponding GEM (BCs or RGCs) was randomly downsampled to include no more than 200 cells per transcriptomic cluster indicated in **Extended Data Figs. 3-6**; (ii) the downsampled species-specific GEMs were combined along the set of shared gene orthologs, normalized to 10,000 counts per cell, and log-transformed; (iii) 2000 highly variable genes were selected within each species, and features that were repeatedly variable were used for anchor finding, integrated dimensionality reduction, and clustering of GEMs based on the Seurat pipeline (Stuart *et al*., 2019). The resulting clusters were called OrthoTypes (OTs). A resolution of 0.5 was used for the clustering. Transcriptomically proximal OTs based on a gene expression dendrogram that contained distinct subsets of species were merged. Note that other than the downsampling step, species cluster IDs were not used to influence the selection of variable genes, integration or clustering steps.

#### Robustness of OTs

The mammalian OTs remained robust to different downsampling trials (data not shown), as well as the inclusion of non-mammals in the analysis (cf. **Fig. 3d-d’’** and **Extended Data Fig. 9d** for BCs, and **Fig. 4d** and **Extended Data Fig. 10c** for RGCs). As the OTs are the result of a clustering of the integrated data, the number of OTs depends on the resolution parameter. We varied the clustering resolution and tracked the number of OTs, the Adjusted Rand Index (ARI) of the clustering, and the number of species-specific OTs. The BC OTs were robust across a wide range of resolution (0.4-1.5), as indicated by a stable number of OTs (16-21), high values of the ARI (0.88-0.96), and very few, if any, species-specific OTs, defined as OTs. The RGC OTs exhibited higher sensitivity to the resolution parameter over the same range, with the number of clusters ranging from 26-46. For resolution values over 1, >5 species-specific OTs were consistently observed across trials. However, ARI values were reasonably high across values tested (0.625-0.849). The results presented in the main text are for a resolution of 0.5.

At early stages of this work, we also repeated the OT analysis using two alternative integration methods – Harmony (Korsunsky et al., 2019) and Liger (Welch et al., 2019). However, the integration with these two methods resulted primarily fragmented, non-specific mapping of individual types within a species. We elected to proceed with Seurat (Stuart *et al*., 2019), given the greater efficacy of its integration while retaining distinctions that reflect “known” biological structure (e.g. ON vs. OFF BCs).

### Factorized Linear Discriminant Analysis (FLDA)

#### Conceptual framework

FLDA is a method that finds a low-dimensional factorization of high-dimensional gene expression data from cells with multiple categorical attributes such that each axis of the low dimensional-space captures the variation along one attribute while minimizing co-variation with other attributes. The mathematical derivations underlying FLDA are described in a previous publication (Qiao and Meister, 2020), and are summarized in **Supplementary Note 3.** In this study, we applied FLDA to factorize transcriptomic data for RGCs carrying three categorical attributes: response polarity (ON vs. OFF), response kinetics (transient vs. sustained), and species (mouse vs. primate). Using A, B, and C to represent these attributes, the total gene expression covariance matrix can be expressed as:

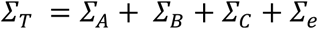

where S_T_ is the total covariance matrix, and S_A_, S_B_, and S_C_ are covariance explained by attributes A, B and C respectively. S_)_ is the residual variance that is not explained by these attributes.

FLDA identifies a 3-D embedding (**u**, **v**, **w**) of the cells such that **u** maximizes the variance of attribute A while minimizing variances of attributes B and C, **v** maximizes the variance of attribute B while minimizing variances of attributes C and A, and **w** maximizes the variance of attribute C while minimizing variances of attributes A and B. **Supplementary Note 3** shows that the **u**, **v** and **w** are solutions to generalized eigenvalue problems.

#### Implementation

To examine the correspondence of primate parasol and midget RGCs with mouse αRGCs, we utilized two published scRNA-seq datasets. The first dataset contains 35,699 adult RGCs, including 399 αRGCs (Tran *et al*., 2019). The second dataset contains 11,724 macaque peripheral RGCs containing 25,399 midget RGCs and 3306 parasol RGCs (Peng *et al*., 2019).

Data was preprocessed and normalized as previously reported (Peng *et al*., 2019; Tran *et al*., 2019). Briefly, transcript counts within each column of the count matrix (genes × cells) were normalized to sum to the median number of transcripts per cell, resulting in normalized counts Transcripts-per-million (*TPM_ij_*) for gene *i* in cell *j*. We used a log-transformed expression matrix *E_ij_ = ln(TPM_ij_ + 1*) for further analysis.

We next identified 7779 high-variance genes (HVGs) using an approach that fits a relationship between the mean and coefficient of variation of gene expression (Chen et al., 2016; Pandey et al., 2018). Next, we performed principal component analysis (PCA) on the dataset to remove multicollinearity. Finally, we analyzed the resulting PCs x cells matrix using FLDA(Qiao and Meister, 2020).

In order to determine the mouse RGC types that best match the four predominant primate RGC types: ON/OFF midgets and ON/OFF parasols, we selected 20 candidates of mouse types with known polarity and kinetics based on previous studies (Goetz et al., 2022; Rousso et al., 2016) (**Supplementary Table 2**). We drew all possible combinations of four RGC types from this set (n=432), and for each combination, we performed FLDA and calculated the eigenvalue corresponding to the polarity and the kinetics axes. We ranked these combinations based on their FLDA eigenvalues and identified the combination with the highest eigenvalue as the best match (**Fig. 5d,d’**).

### Code Availability

scRNA-seq data clustering, integration, and visualization was performed in the R statistical language, and heavily relied on the Seurat package (https://satijalab.org/seurat/). All scripts are available at https://github.com/shekharlab/RetinaEvolution including R markdown notebooks. FLDA analysis was performed in Python, and the code and documentation are available at https://github.com/muqiao0626/FLDA.

## Extended data figures and tables

**Extended Data Fig. 1.**
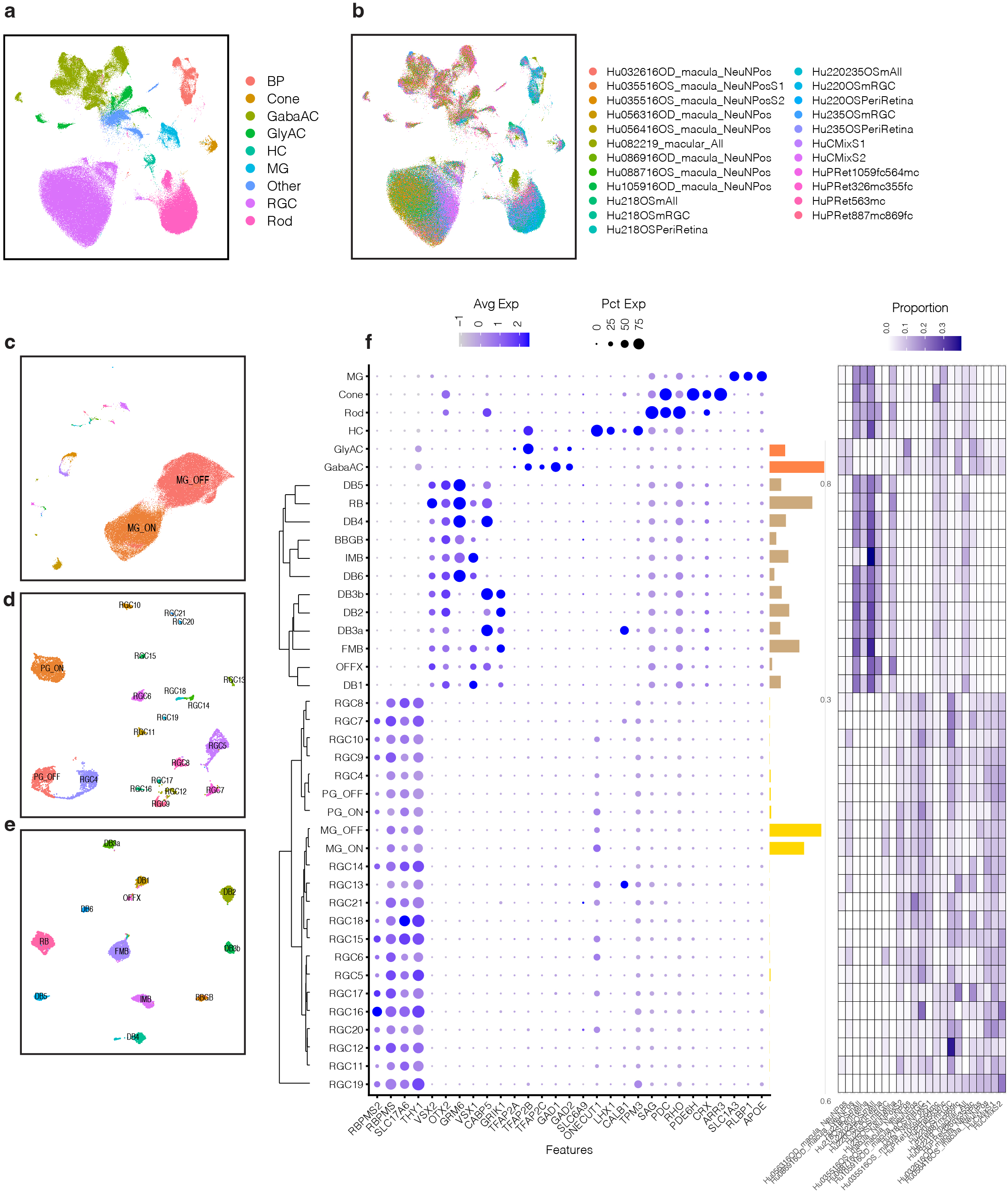
snRNA-seq data from the fovea/macula and peripheral retina of healthy human donors. a. UMAP embedding of 184,808 nuclei from the central and peripheral retina of healthy human donors, with individual points colored by cell class. PRs have been divided into rod and cone subclasses, and ACs have been divided into GABAergic and glycinergic subclasses. b. Same as **a**, with points colored by sample identity. c. UMAP embedding of 80,032 RGC nuclei from the foveal and peripheral retina of healthy human donors, with individual points colored by type identity. Only ON and OFF midget ganglion RGCs are labeled. d. UMAP embedding of 6615 non-midget RGC nuclei from **c**, with individual points colored by type identity. ON and OFF parasol ganglion cells are labeled. e. UMAP embedding of 9126 BC nuclei from the fovea and peripheral retina of healthy human donors, with individual points colored by type identity. f. Dotplot showing expression of cell class-specific markers (columns) in the human clusters (rows). The size of each dot represents the fraction of cells in the group with non-zero expression, and the color represents expression level. The six classes are MG, HC, PR (subdivided into Rod and Cone), AC (subdivided into Gabaergic ACs (GabaAC) and glycinergic ACs (Gly AC)), BC and RGC. Only BCs and RGCs have been subclustered. Rows corresponding to BC and RGC clusters are ordered based on hierarchical clustering (dendrograms, left). Barplot on the right of the dotplot depicts the relative frequency of each cluster within a class (colors). The rightmost heatmap depicts the distribution of each cluster across biological replicates (columns).

**Extended Data Fig. 2.**
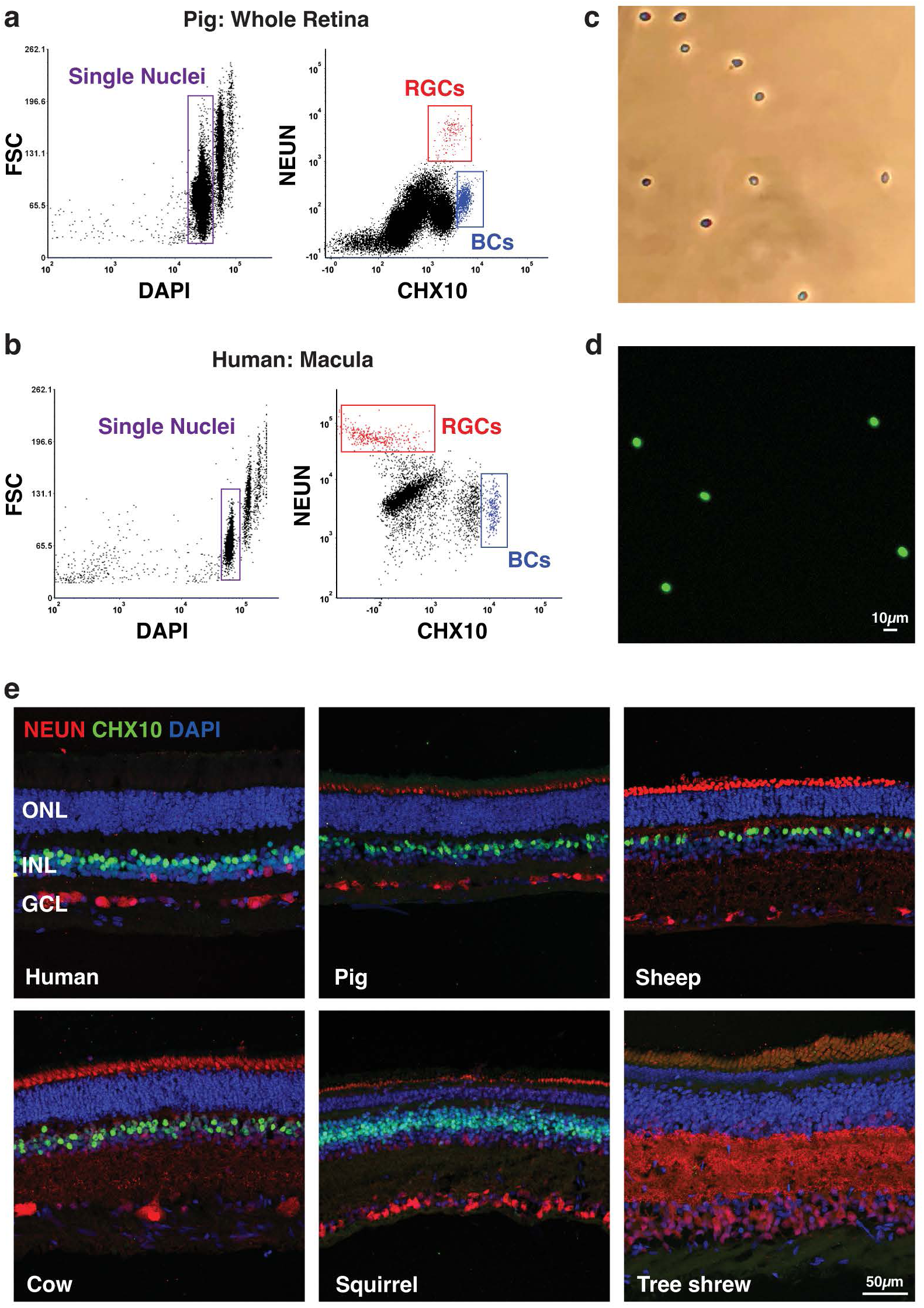
Nuclear enrichment strategies for retinal ganglion cells (RGCs) and bipolar cells (BCs). a. Examples of gating strategy in fluorescent activated cell sorting (FACS) experiments for collecting single nuclei labeled with either PE-conjugated NEUN, which enriches RGCs, or APC-conjugated CHX10 (also known as VSX2), which enriches BCs. Data shown are from experiments in the pig retina. NEUN and CHX10-based enrichment resulted in ∼90% yield for RGCs and ∼95% yield for BCs. b. Same as panel **a**, for human macular retina samples. NEUN-based enrichment resulted in ∼90% yield for RGCs; BCs were not analyzed in this experiment. c. Brightfield image showing the morphology and integrity of FACS-purified nuclei. d. Confocal image of DAPI stained FACS-purified nuclei. e. Retinal sections from six species show that PE-conjugated NEUN (red) and APC-conjugated CHX10/VSX2 labels RGCs and BCs, respectively. Retinal sections were co-stained for DAPI (blue) to visualize nuclei. Scale bar, 50 µm.

**Extended Data Fig. 3.**
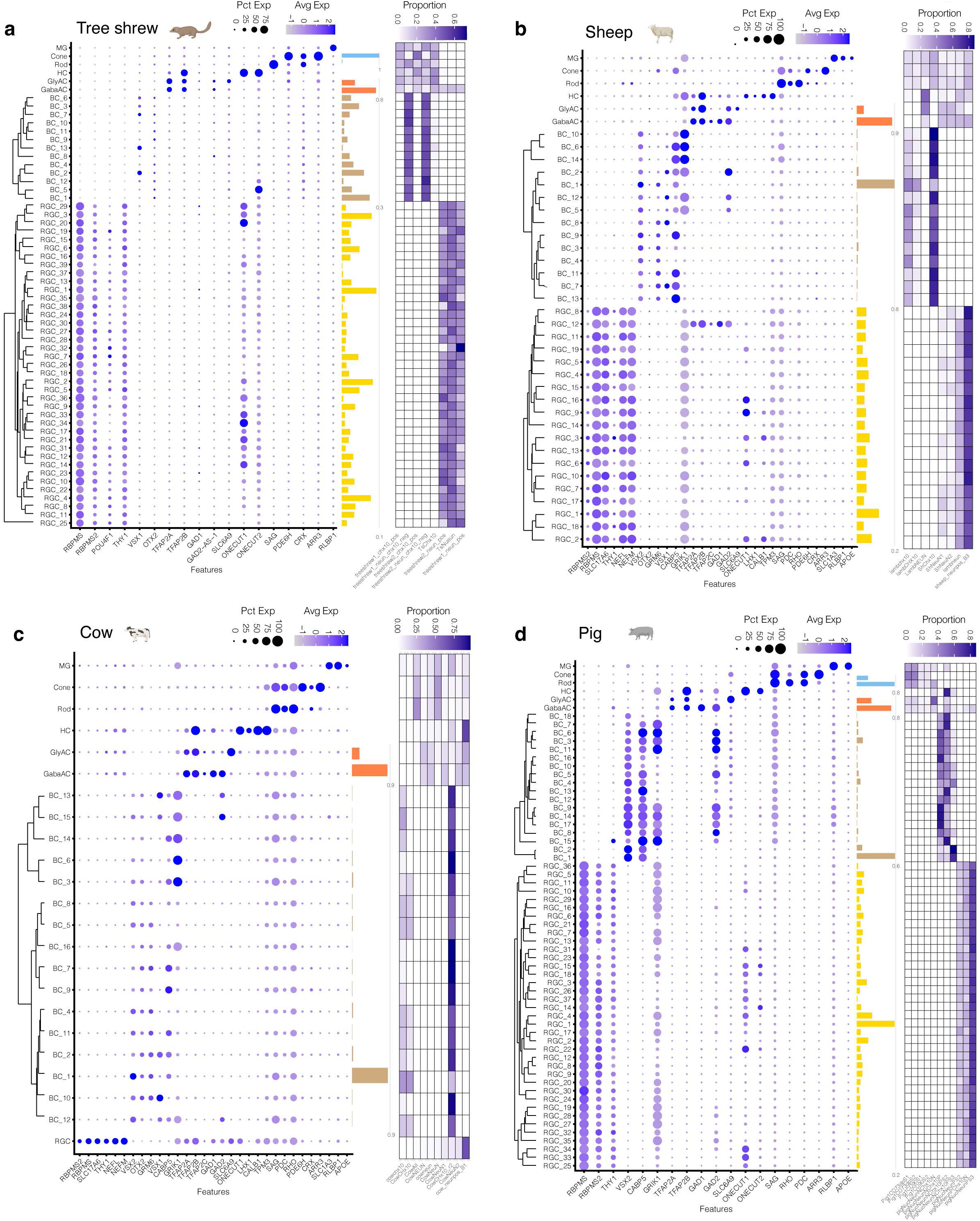
Summary of cell type atlases for tree shrew, sheep, cow, and pig. a. Dotplot showing expression of cell class-specific markers (columns) in the tree shrew clusters (rows). The size of each dot represents the fraction of cells in the group with non-zero expression, and the color represents expression level. The six classes are MG, HC, PR (subdivided into Rod and Cone), AC (subdivided into GABAergic AC (GabaAC) and glycinergic AC (Gly AC)), BCs and RGCs. Only BCs and RGCs have been subclassified through a within-species integration and clustering analysis (**Methods**). Rows corresponding to BC and RGC clusters are ordered based on a hierarchical clustering analysis (dendrograms, left). Barplot on the right of the dotplot depicts the relative frequency of each cluster within a class (colors). The rightmost heatmap depicts the distribution of each cluster across biological replicates (columns). Panels **b-d** depict the same information as panel **a** for sheep (**b**), cow (**c**), and pig (**d**).

**Extended Data Fig. 4.**
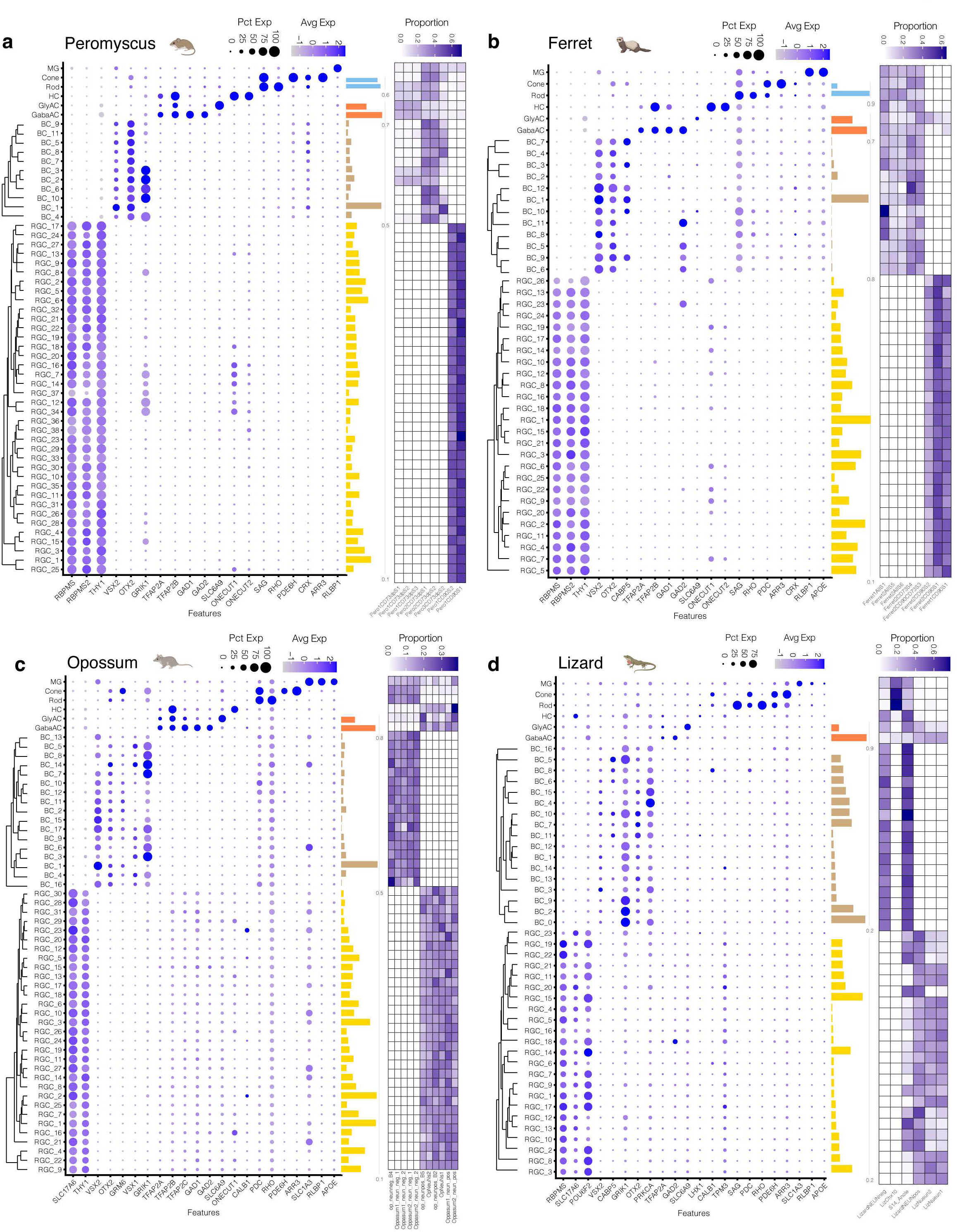
Summary of cell type atlases for Peromyscus, ferret, opossum, and brown anolis lizard. Panels **a-d** depict the atlases (as in **Extended Data Fig. 3**) for peromyscus (**a**), ferret (**b**), opossum (**c**), and brown anolis lizard (**d**).

**Extended Data Fig. 5.**
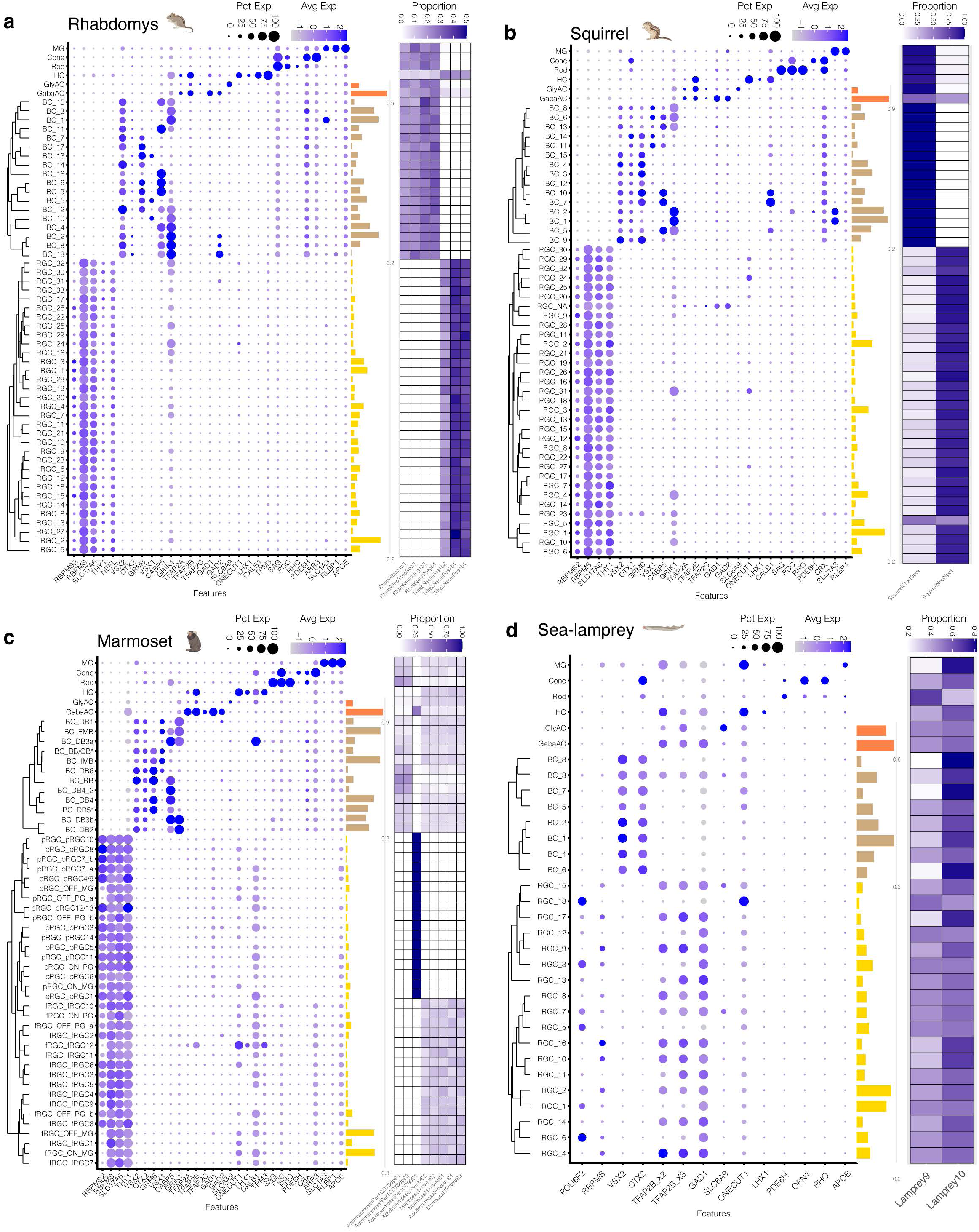
Summary of cell type atlases for Rhabdomys, squirrel, marmoset and sea-lamprey. Panels **a-d** depict atlases (as in **Extended Data Fig. 3**) for rhabdomys (**a**), squirrel (**b**), marmoset (**c**), and Sea-lamprey (**d**).

**Extended Data Fig. 6.**
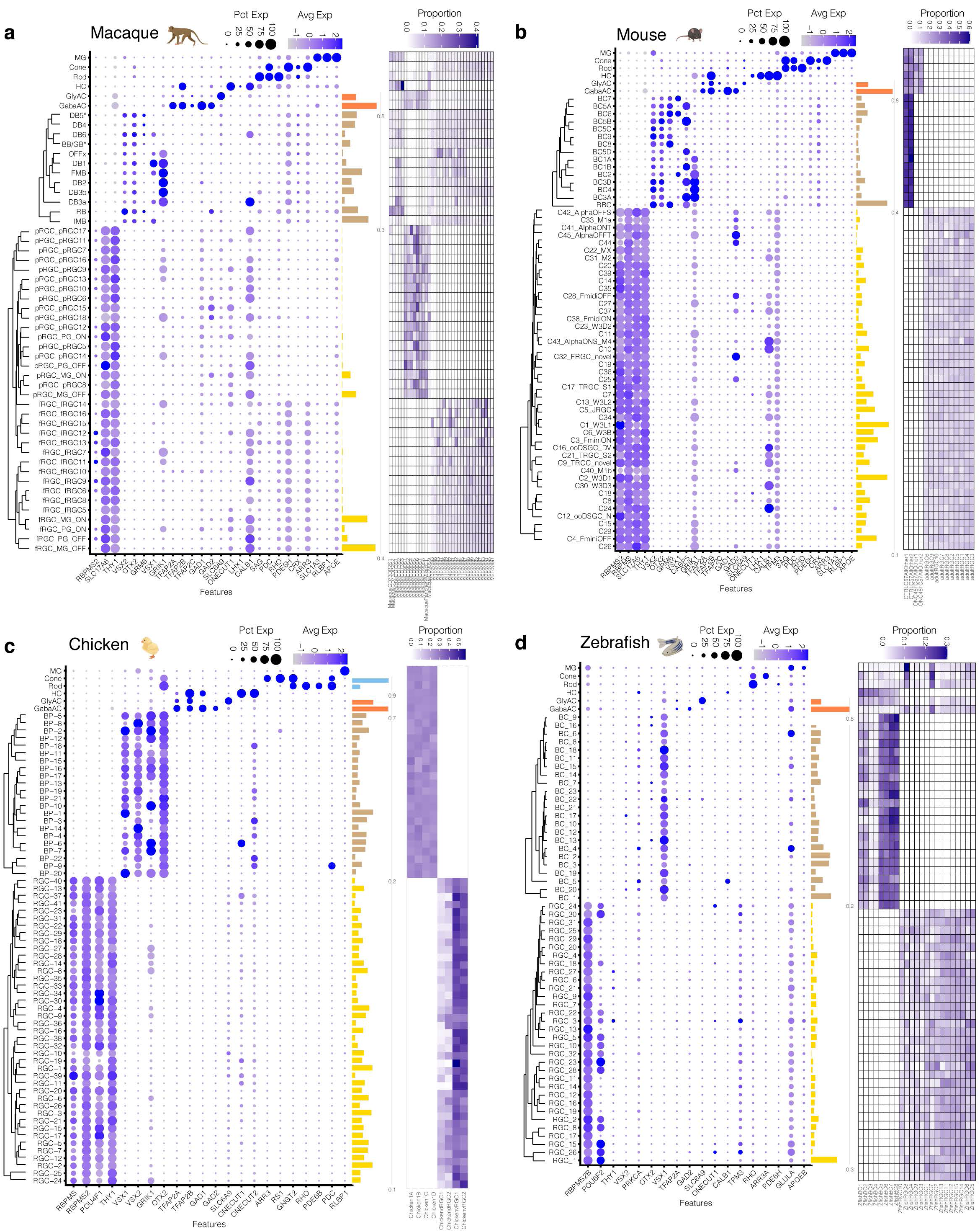
Summary of cell type atlases for macaque, mouse, chick and zebrafish. Panels **a-d** depict atlases (as in **Extended Data Fig. 3**) for macaque (**a**), mouse (**b**), chick (**c**), and zebrafish (**d**). Cluster labels are consistent with published annotations (Kolsch, *et al*., 2021; Peng *et al*., 2019; Shekhar *et al*., 2016; Tran *et al*., 2019; Yamagata *et al*., 2021).

**Extended Data Fig. 7.**
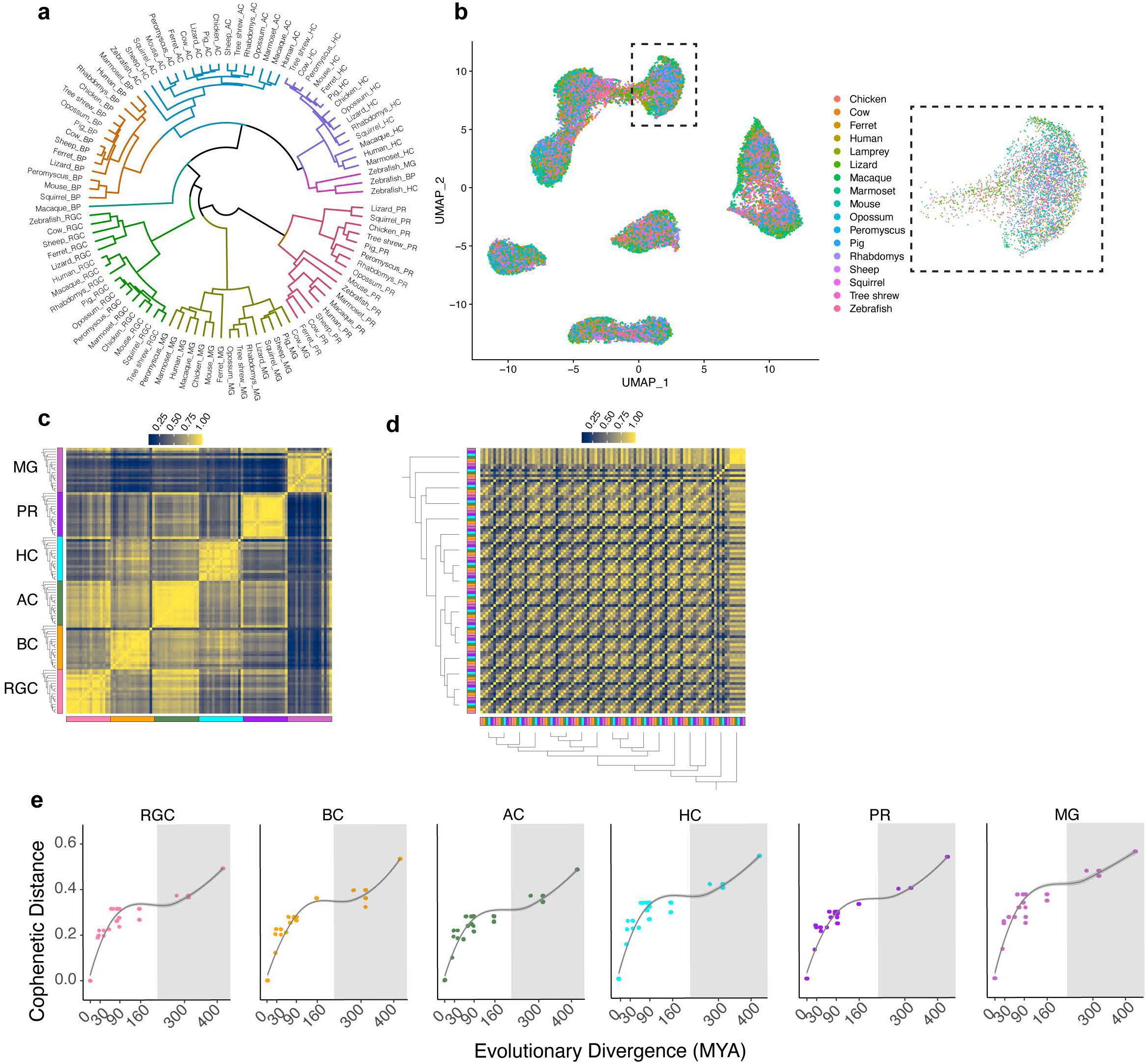
Evolutionary conservation of retinal classes. a. Dendrogram showing transcriptional relationships among pseudobulk expression vectors following integration. Each node is a cell class within a particular species. Dendrograms were computed via hierarchical clustering analysis (correlation distance, average linkage). b. Same as **Fig. 2d**, with cells colored by species of origin. Inset shows a magnified region containing samples from all species. c. Cross-correlation matrix (spearman) of class- and species-specific cell-averaged profiles for all 17 vertebrates (compare with **Fig. 2b**). Rows and columns are grouped by class, and then ordered by phylogeny within a class. d. Same as panel c, but rows and columns grouped based on species instead of class (compare with **Fig. 2c**). e. Pairwise cophenetic distance (y-axis) within gene expression dendrograms is correlated with evolutionary distance (x-axis). The dendrograms were computed for each class separately via hierarchical clustering (correlation distance, average linkage) (**Methods**). The cophenetic distance between two nodes is the height of the dendrogram where the two branches that include the two objects merge into a single branch. Panels correspond to different classes.

**Extended Data Fig. 8.**
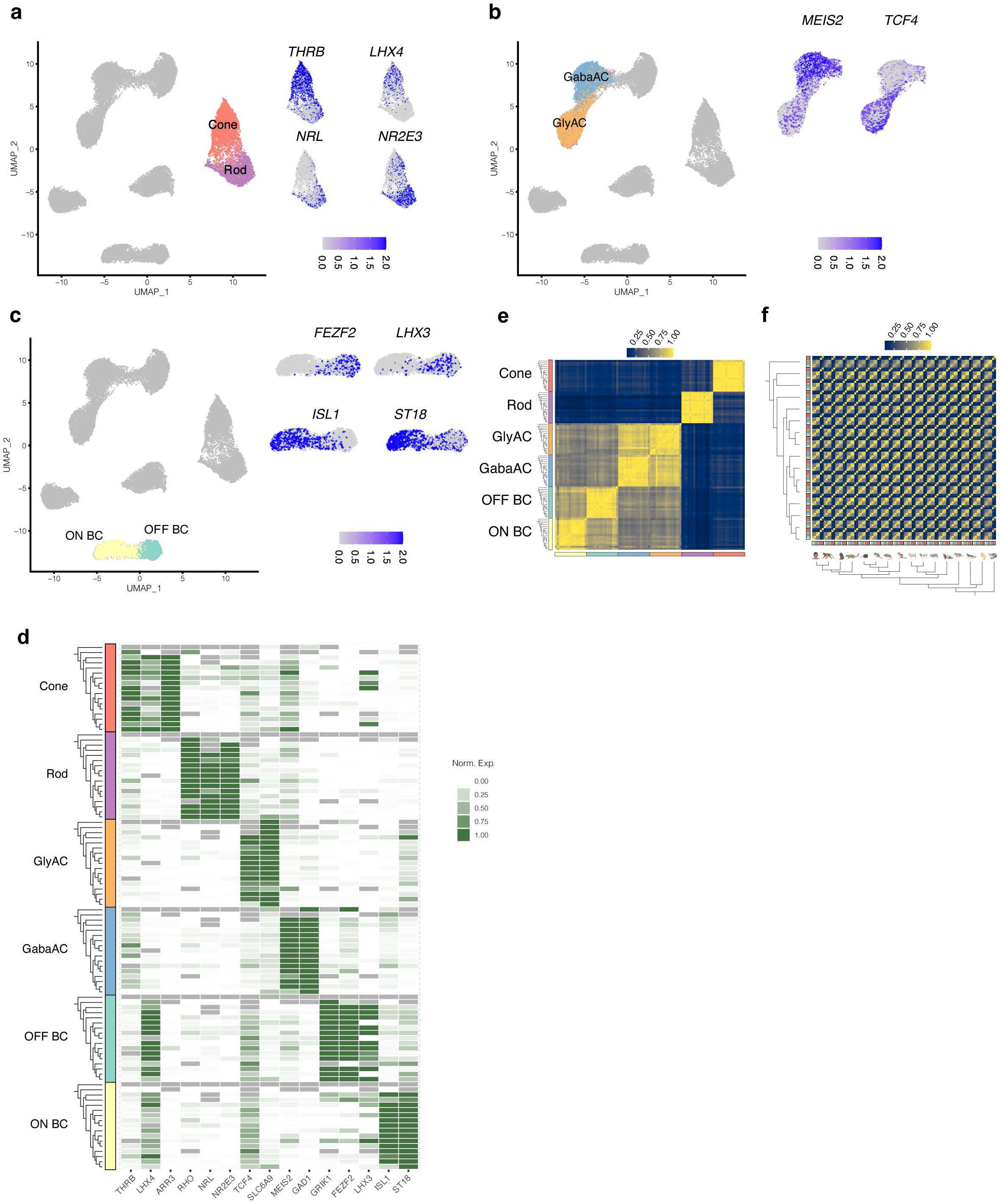
Evolutionary conservation of retinal subclasses. a. UMAP embedding of integrated cross-species data (as in Fig. 2d), highlighting PR subclasses cones and rods. Insets show feature plots of cone-specific (top) and rod-specific (bottom) transcription factors (TFs). b. Same as panel **a**, for AC subclasses GABAergic ACs (GabaAC) and glycinergic ACs (GlyAC). Insets show feature plots of a GABAergic TF *MEIS2* and a glycinergic TF *TCF4*. c. Same as panel **a**, for BC subclasses ON BCs and OFF BCs. Insets show feature plots of OFF BC-specific (top) and ON BC-specific (bottom) transcription factors (TFs). d. Heatmap showing average expression of subclass-specific genes (columns) within the six subclasses across 17 species (rows). Rows are grouped by subclass (annotation bar, left). Within each subclass, species are ordered as in **Fig. 1b**, with top and bottom nodes in each dendrogram corresponding to lamprey and human, respectively (corresponding to right and left in **Fig. 1a**). e. Cross-correlation matrix (spearman) of subclass- and species-specific pseudobulk transcriptomic profiles for all 16 jawed vertebrates. Rows and columns are grouped by subclass, and then ordered by phylogeny within a class. Lamprey was excluded due to paucity of shared orthologs. f. Same as panel **d**, but rows and columns grouped based on species instead of subclass.

**Extended Data Fig. 9.**
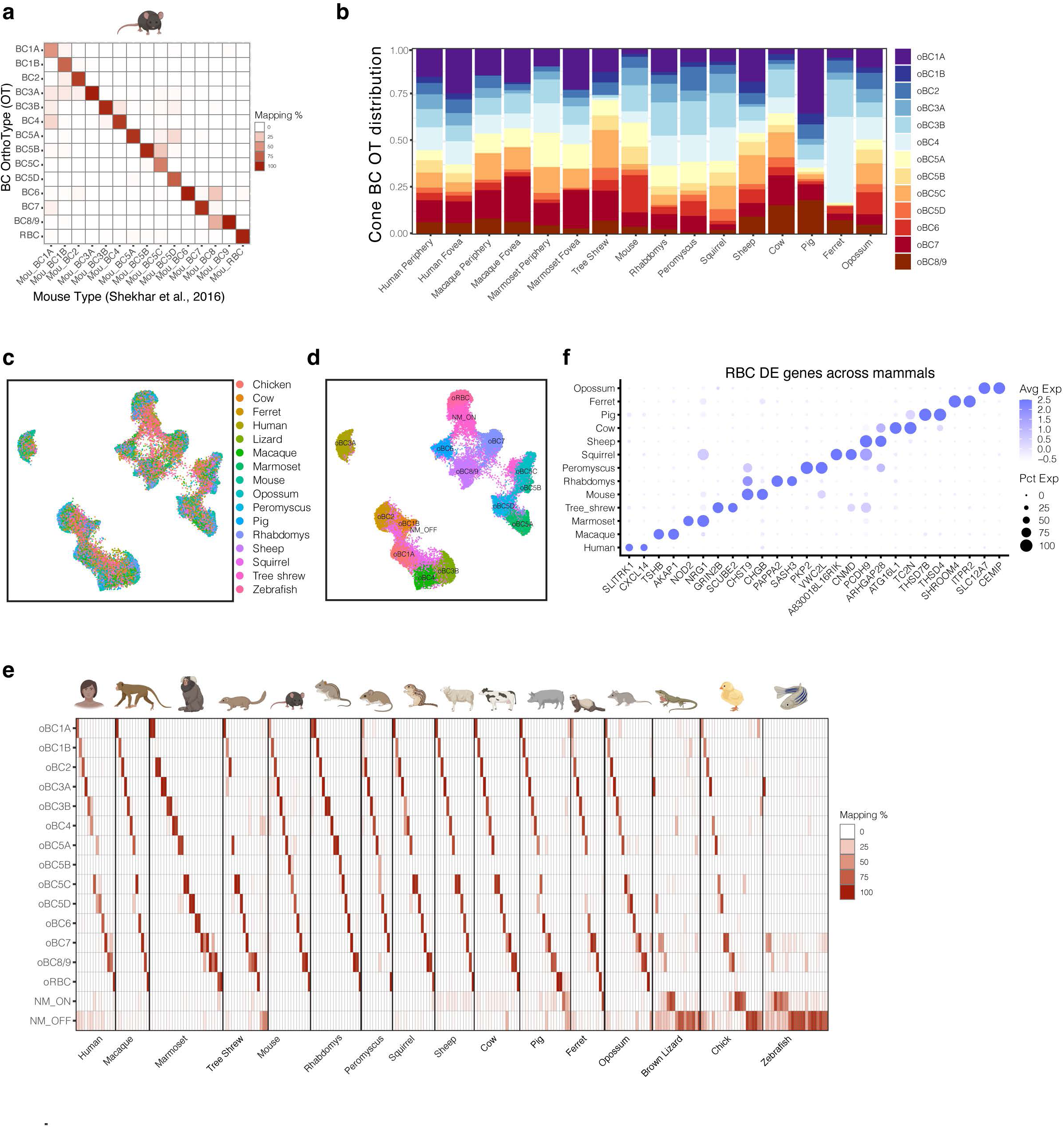
Bipolar Cell OrthoType analysis including non-mammals. a. Confusion matrix showing the rationale behind naming mammalian BC OTs (rows) based on the mapping patterns of mouse BC types (columns) (Shekhar, et al., 2016). Representation as in **Fig. 3d**, with each column summing to 100%. OT BC8/9 contains mappings from both mouse BC8 and BC9, which are transcriptionally proximal. b. Barplot showing within-species relative frequencies (y-axis) of the 13 cone BC OTs within each mammalian species (x-axis). The foveal and peripheral data from primates are plotted separately. c. Integrated UMAP of BCs from all 16 jawed vertebrates. Cells are colored by species of origin. Lamprey, a jawless vertebrate, was excluded from the analysis due to the paucity of shared orthologous genes. d. Same as **c**, with cells colored by OT identity. The integration of all jawed vertebrates recovers all the mammalian BC OTs listed in **Fig. 3c**, but additionally identifies two OTs enriched for non-mammalian BCs from chick, lizard and zebrafish. The two OTs, named NM_OFF and NM_ON, are enriched for OFF and ON BCs from non-mammals (also see panel **e**). e. Confusion matrices showing the mapping of species-specific BC clusters (columns) to BC OTs (rows) identified by integrating BCs from all jawed vertebrates (panel c). Representation as in **Fig. 3d’**. Mammalian BC clusters predominantly map to the mammalian OTs (rows 1-14), and the pattern of mapping is similar to **Fig. 3d**. Chick, Lizard and Zebrafish BCs largely map to the non-mammalian OTs NM_OFF and NM_ON (rows 15-16). f. Dotplot showing species-specific genes (columns) expressed in RBC orthologs in mammals (rows). The size and color of each dot represent the percentage of cells within the species cluster expressing the gene and the average expression level, respectively.

**Extended Data Fig. 10.**
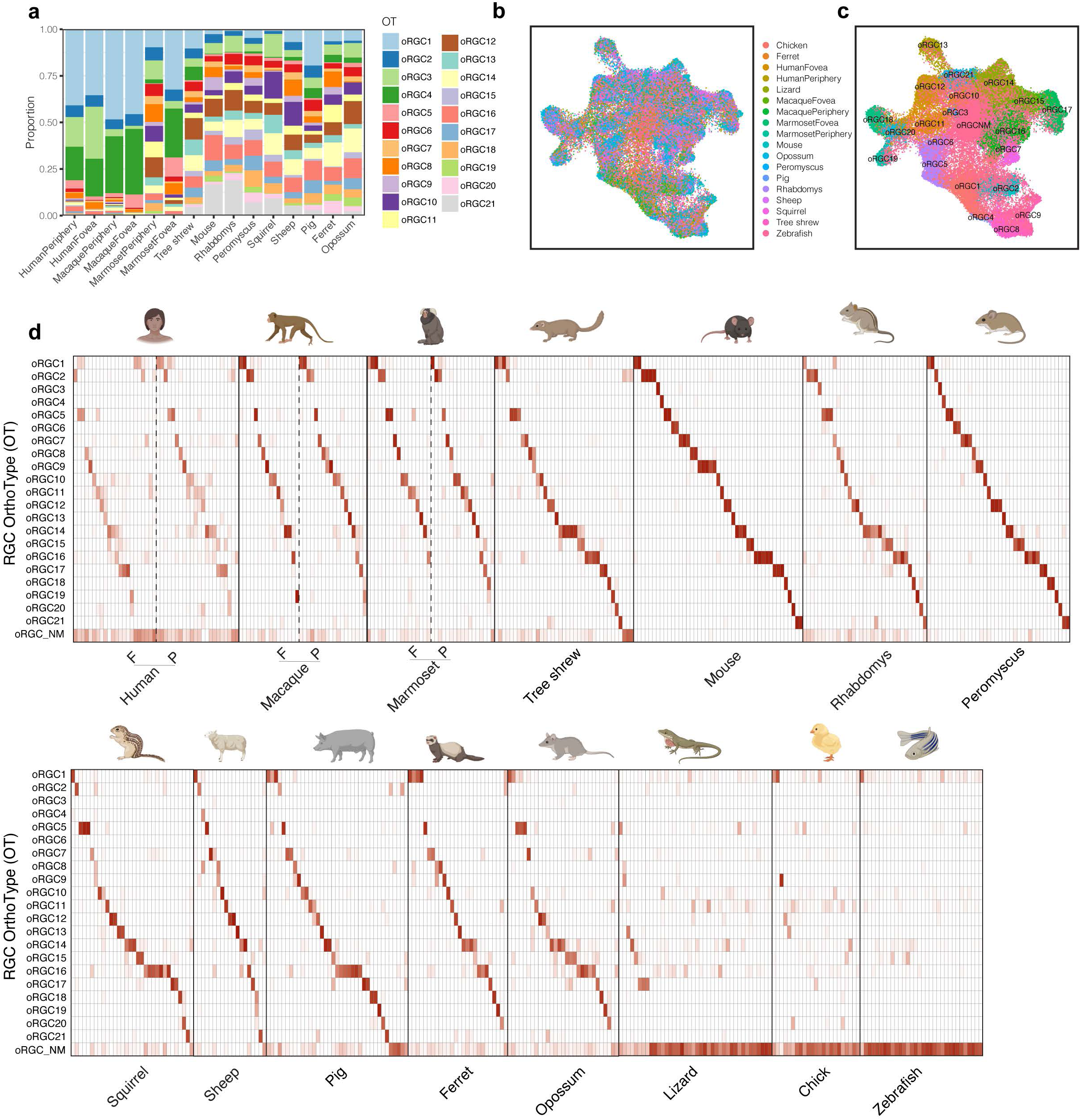
Retinal Ganglion Cell OrthoType analysis including non-mammals. a. Barplot showing within-species relative frequencies (y-axis) of the 21 RGC OTs within mammalian species (x-axis) (**Fig. 4b**). The foveal and peripheral data from primates are shown separately. b. Integrated UMAP of RGCs from all 15 jawed vertebrates (excluding cow). Cells are colored by species of origin. For primates, fovea and periphery are plotted separately. c. Same as **b**, with cells colored by RGC OT. OTs 1-21 map 1:1 to the mammalian OTs in **Fig. 4b**, but we recover an additional OT (NM) predominantly containing non-mammalian RGCs from chick, lizard and zebrafish (also see panel **d**). d. Confusion matrices showing the mapping of species RGC clusters (columns) to RGC OTs (rows) identified by integrating RGCs from all jawed vertebrates (panel c). Representation as in **Fig. 4d**. Mammalian RGC clusters predominantly map to the mammalian OTs (rows 1-21), and the pattern of mapping is similar to **Fig. 4d**. With the exception of ipRGCs, chick, lizard and zebrafish RGCs largely map the NM OT (row 22).

**Extended Data Fig. 11.**
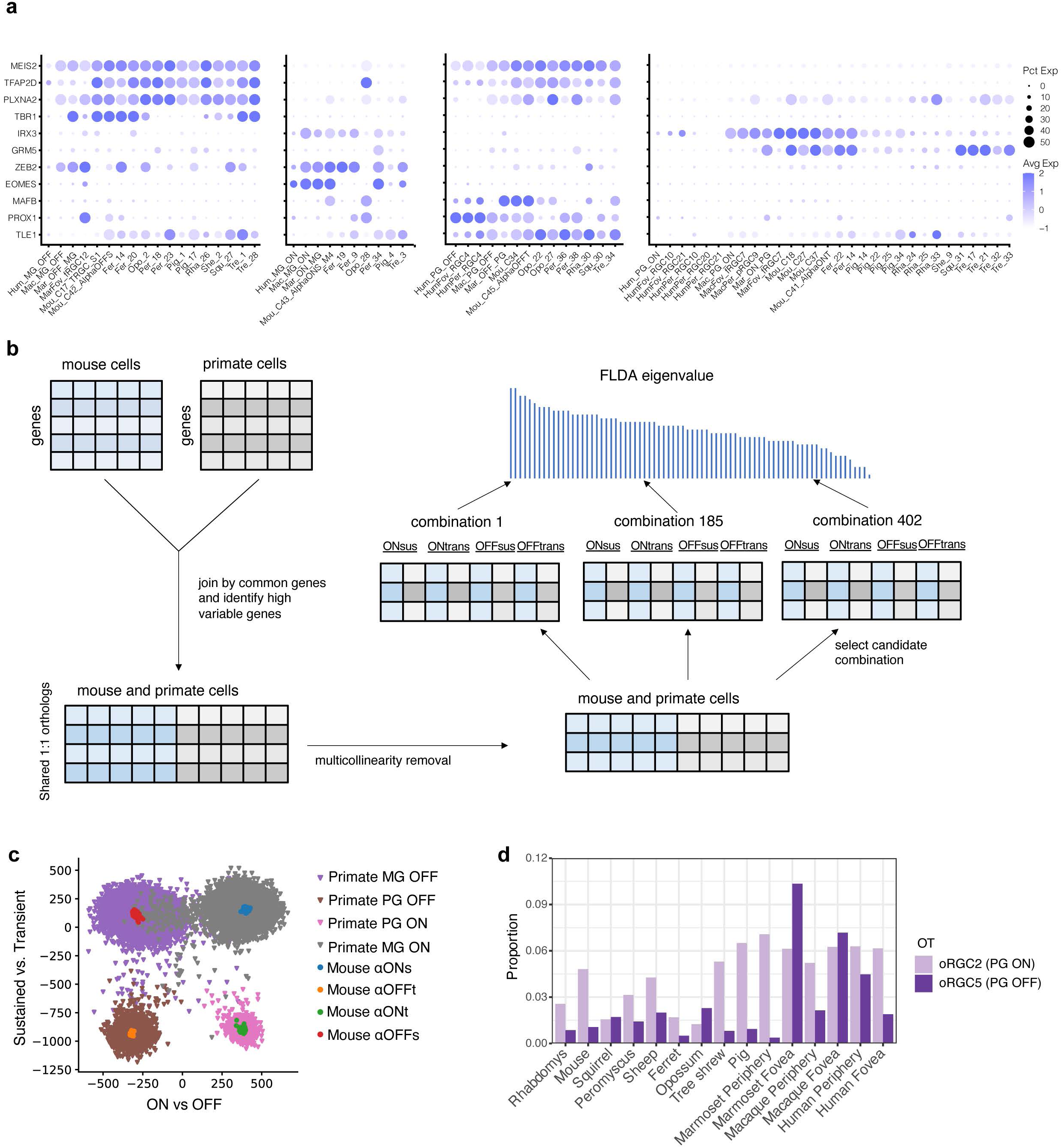
Midget and Parasol OTs. a. Dotplot showing examples of DE genes across OT1-4 and their expression across orthologous species-specific clusters. The size and color of each dot represent the percentage of cells within the species cluster expressing the gene and the average expression level, respectively. Column order as in **Fig. 5a**. b. FLDA workflow and eigenvalue analysis. The gene expression matrices of primate and mouse RGCs were combined by their shared orthologous genes. Highly variable genes were selected, and PCA was applied to remove multicollinearity. FLDA was performed on different combinations of mouse RGC candidates with known polarity and kinetics listed **Supplementary Table 2**. The combinations were ranked based on their FLDA eigenvalues, which measures the variance along each attribute captured in the projection. c. Visualization of the FLDA projection (**Fig. 5c**) along the 2D subspace corresponding to polarity (x-axis) and kinetics (y-axis) d. Relative proportion of parasol RGC orthologs in mammalian species based on the frequencies of cells in oRGC2 and oRGC5.

## Supplementary Note 1

**Fig. 3g** plots rods as a proportion of all PRs and RBCs as a proportion of all BCs for species considered here. RBC proportions were calculated from the respective scRNA-seq and snRNA-seq datasets (**Extended Data Fig. 3-6**). The proportions of rods were obtained from previous reports (see **Table A** below). Only for the case of lizards, we calculated rod proportions from PRs in our data, as we could not obtain any published estimates.

Within the fovea of humans, macaques, and marmosets, rod proportions were calculated by dividing the estimated density of rods to the sum of the density of rods and cones densities. The rod and cone densities were calculated from the original reports by averaging values at five eccentricities (up to 0.5mm) (ref. 1: Fig. 7 and Fig. 9A; ref. 2: Fig. 2; ref. 5: Fig. 3; ref. 6: Fig. 15)

**Table A:** Rod proportions by species and literature sources

| Species/Region | Scientific name | Rod % | Ref |
| --- | --- | --- | --- |
| Human total | *Homo sapiens* | 95.24 | 1,2,3 |
| Macaque | *Macaca fascicularis* | 95.05 | 2,4,5 |
| Marmoset | *Callithrix jacchus* | 83.33 | 6,7 |
| Tree shrew | *Tupaia belangeri chinensis* | 8.86 | 8 |
| Pig | *Sus domesticus* | 88.89 | 9 |
| Sheep | *Ovis aries* | 91.67 | 10 |
| Cow | *Bos taurus* | 92.00 | 11 |
| Cpossum | *Monodelphis domestica* | 99.00 | 12 |
| Ferret | *Mustela putoriusfuro* | 93.20 | 13 |
| Rhabdomys | *Rhabdomys pumilio* | 55.16 | 14 |
| Peromyscus | *Peromyscus maniculatus bairdii* | 98.50 | 15 |
| Mouse | *Mus musculus* | 97.26 | 16,17 |
| Squirrel | *Ictidomys tridecemlineatus* | 14.48 | 18,19 |
| Zebrafish (adult) | *Danio rerio* | 47.00 | 20 |
| Chicken | *Gallus gallus domesticus* | 40.00 | 21 |
| Lizard | *Anolis sagrei* | 1.00 | our data |
| Human 1mm fovea (estimate) | *Homo sapiens* | 16.00 | 1,2,3 |
| Macaque 1mm fovea (estimate) | *Macaca fascicularis* | 16.00 | 2,4,5 |
| Marmoset 1mm fovea (estimate) | *Callithrix jacchus* | 21.00 | 6,7 |

## Supplementary Note 2

Approximate morphologies and stratification of four types of cone bipolar cells and four types of RGCs in mouse and primate retinas were drawn based on the illustrations from previous literature listed in the table below.

**Table 1:** A list of figures in previous reports that were used to draw rough morphology of bipolar cells and RGCs depicted in Fig. 5b.

| Class | Type | Species | Source |
| --- | --- | --- | --- |
| BC | BC1, BC7, BC3a, BC5, FMB, IMB, DB3a, DB4 | *M. musculus, M. fascicularis* | Fig. 8 in Ref. 1 |
| BC | BC1, BC7, BC3a, BC5 | *M. musculus* | Fig. 1b in Ref. 2 |
| BC | BC1, BC7, BC3a, BC5 | *M. musculus* | Fig. 2 in Ref. 3 |
| BC | FMB, IMB, DB3a, DB4 | *M. fascicularis* | Fig. 1 in Ref. 4 |
| RGC | ON Midget, ON Parasol | *M. fascicularis, H sapiens* | Fig. 9 in Ref. 4 |
| RGC | ON Parasol, OFF Parasol, OFF midget | *M. fascicularis, H sapiens* | Fig. 4a in Ref. 5 |
| RGC | α-OFFt | *M. musculus* | Fig. 2a in Ref. 6 |
| RGC | α-OFFs, α-ONs, α-OFFt | *M. musculus* | Fig. 3 in Ref. 7 |
| RGC | α-OFFs, α-ONs, α-OFFt, α-ONt | *M. musculus* | Fig. 4b in Ref. 8 |

## Supplementary Note 3

For a detailed description of the FLDA method, please refer to our previous work (1). In brief, FLDA is a method that projects high-dimensional gene expression data from cells with multiple categorical attributes into a low-dimensional space where each axis captures the variation along one attribute while minimizing co-variation with other attributes.

In this study, we used FLDA to analyze three categorical attributes of retinal neurons: response polarity (ON vs. OFF), response kinetics (transient vs. sustained), and species (mouse vs. primate). Let’s use A, B, and C to represent these attributes. *i*, *j*, *k* denote the indices of attributes A, B, and C, and *a*, *b*, *c* are the number of categories in attributes A, B, and C. *n_ijk_*is the number of cells in the category combination *ijk*. ***x****_ijkl_* is the gene expression vector of the *l*th cell in the category combination *ijk*.

The covariance matrix of total variance can be decomposed as:

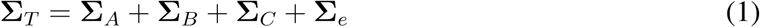

where

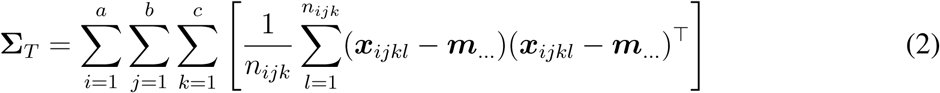

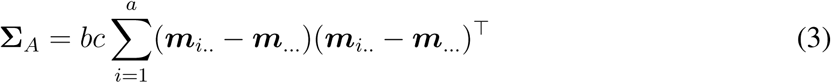

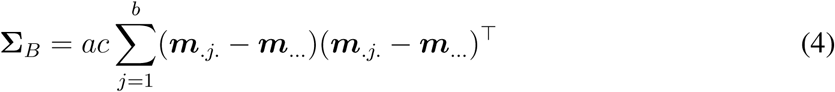

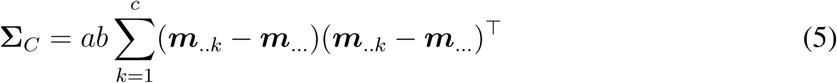

and

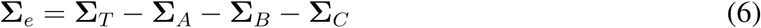

**Σ***_T_* is the total covariance matrix, and **Σ***_A_*, **Σ***_B_*, and **Σ***_C_* are covariance explained by attributes A, B and C respectively. **Σ***_e_* is the residual variance that is not explained by these attributes.

Here,

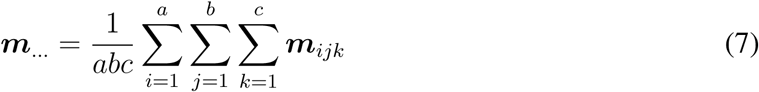

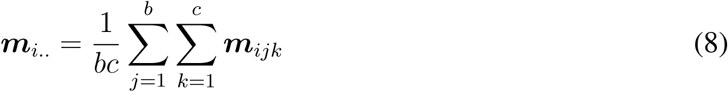

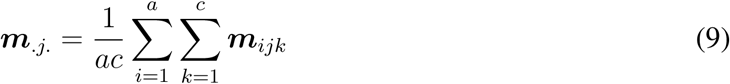

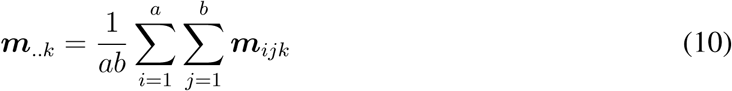

and

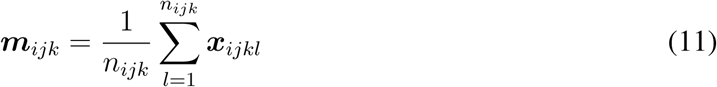

Our objective was to find a projection that maximizes the variance of attribute A while minimizing the variances of attributes B and C. Specifically, we aimed to find *u^∗^* that maximizes the following equation:

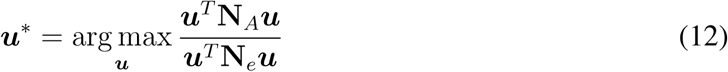

where

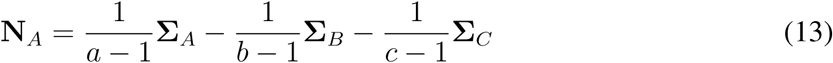

and

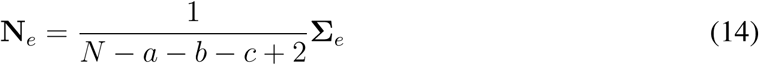

*N* is the total number of cells, and *a −* 1, *b −* 1, *c −* 1, and *N − a − b − c* + 2 are the degrees of freedom of the corresponding terms.

This optimization problem is commonly referred to as a generalized eigenvalue problem (2). Here, **N***_A_* is symmetric but not necessarily positive definite, and **N***_e_* is positive definite. When **N***_e_* is invertible, the eigenvector *u^∗^* associated with the largest eigenvalue of 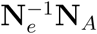 is selected. In this study, we identify the eigenvector with the largest eigenvalue of 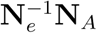, which we refer to as the FLDA eigenvalue for the attribute A. This FLDA eigenvalue measures how much variance of the corresponding attribute (A) is captured compared to the variances of other attributes (B, C). The eigenvector *u^∗^* can be normalized to have a unit length. The elements within the unit vector represent the relative weights of the corresponding genes.

Similarly, to find a low-dimensional representation aligned with a categorical attribute *B*, we maximized the objective:

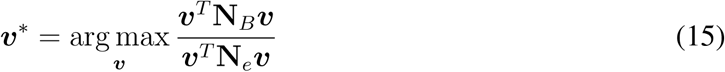

where

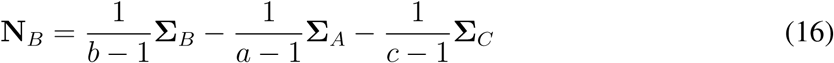

and to find a low-dimensional representation aligned with a categorical attribute *C*, we maximized the objective:

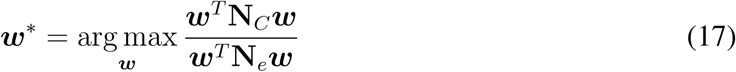

where

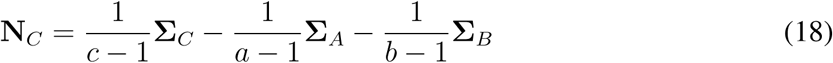

